# CCDC32 stabilizes clathrin-coated pits and drives their invagination

**DOI:** 10.1101/2024.06.26.600785

**Authors:** Ziyan Yang, Changsong Yang, Zheng Huang, Peiliu Xu, Yueping Li, Lu Han, Linyuan Peng, Xiangying Wei, John Pak, Tatyana Svitkina, Sandra L. Schmid, Zhiming Chen

**Affiliations:** NHC Key Laboratory of Birth Defect Research and Prevention, MOE Key Laboratory of Rare Pediatric Diseases, Institute of Cytology and Genetics of School of Basic Medical Sciences & Department of Clinical Laboratory of The First Affiliated Hospital, Hengyang Medical School, University of South China, Hengyang, Hunan, China; Department of Biology, University of Pennsylvania, Philadelphia, PA, USA; Fuzhou Institute of Oceanography, College of Geography and Oceanography, Minjiang University, Fuzhou, Fujian, China; Chan Zuckerberg Biohub, San Francisco, California, USA; Department of Cell Biology, University of Texas Southwestern Medical Center, Dallas, Texas, USA

## Abstract

Clathrin-mediated endocytosis (CME) is essential for maintaining cellular homeostasis. Previous studies have reported more than 50 CME accessory proteins; however, the mechanism driving the invagination of clathrin-coated pits (CCPs) remains elusive. We show by quantitative live cell imaging that siRNA-mediated knockdown of CCDC32, a poorly characterized endocytic accessory protein, leads to the accumulation of unstable flat clathrin assemblies. CCDC32 interacts with the α-appendage domain (AD) of AP2 *in vitro* and with full length AP2 complexes in cells. Deletion of aa78-98 in CCDC32, corresponding to a predicted α-helix, abrogates AP2 binding and CCDC32’s early function in CME. Furthermore, clinically observed nonsense mutations in CCDC32, which result in C-terminal truncations that lack aa78-98, are linked to the development of cardio-facio-neuro-developmental syndrome (CFNDS). Overall, our data demonstrate the function of a novel endocytic accessory protein, CCDC32, in regulating CCP stabilization and invagination, critical early stages of CME.

**Summary:** We show that CCDC32, a poorly studied and functionally ambiguous protein, binds to AP2 and regulates CCP stabilization and invagination. Clinically observed mutations in CCDC32 lose their ability to interact with AP2 likely contributing to the development of cardio-facio-neuro-developmental syndrome.

## Introduction

Clathrin-mediated endocytosis (CME) regulates nutrient uptake and maintains the activity of transmembrane transporters and is thus, essential for maintaining cellular homeostasis (Kaksonen and Roux, 2018; Kirchhausen et al., 2014; McMahon and Boucrot, 2011; Mettlen et al., 2018). Malfunctions of CME are strongly associated with neurological diseases, cardiovascular diseases, and cancers (Blue et al., 2018; DeMari et al., 2016; Elkin et al., 2015; Gilles Moulay, 2019; Hamdan et al., 2017; Manti et al., 2019; Sznajder and Swanson, 2019; Wu and Yao, 2009). CME occurs via the assembly of clathrin triskelia into clathrin-coated pits (CCPs) that invaginate to form clathrin-coated vesicles (CCVs) (Kaksonen and Roux, 2018; Mettlen et al., 2018; Smith and Smith, 2022). During CCP maturation, successful invagination of the clathrin coat and its underlying membrane is a key step that determines whether nascent CCPs are productive or abortive (Aguet et al., 2013; Baschieri et al., 2020; Chen and Schmid, 2020; Wang et al., 2020). Although more than 50 endocytic accessory proteins (EAPs) have been reported to be involved in the progression of CME (Bhave et al., 2020; Chen and Schmid, 2020; Kirchhausen et al., 2014; McMahon and Boucrot, 2011; Merrifield and Kaksonen, 2014; Sochacki et al., 2017; Taylor et al., 2011; Traub, 2011), the mechanism of CCP invagination remains elusive, i.e. which accessory proteins are required and how do they regulate CCP invagination?

Coiled-coil domain-containing protein 32 (CCDC32), also known as gene *C15orf57*, is a small, 185 amino acids (aa) protein that has been poorly studied and whose function was unknown. Clinical genome sequencing of three patients with cardio-facio-neuro-developmental syndrome (CFNDS) revealed three homozygous nonsense mutations that only express the first 9, 54 and 80aa of CCDC32, respectively (Abdalla et al., 2022; Harel et al., 2020). How these loss-of-function mutations result in CFNDS remains unknown.

A genome-wide co-essential modules study (Wainberg et al., 2021) suggested a functional correlation between CCDC32 and AP2, whose interactions were also implied by affinity purification-mass spectroscopy (AP-MS) analysis (Cho et al., 2022) and co-immunoprecipitation (co-IP) assays (Wainberg et al., 2021). In addition, depletion of CCDC32 was observed to inhibit transferrin uptake, suggesting a role in CME (Wainberg et al., 2021). More recently, and while this paper was under review, Wan et al. (Wan et al., 2024) reported an essential role for CCDC32 as a co-chaperone with AAGAB (alpha and gamma adaptin binding protein) in the assembly of AP2 complexes. The authors showed that in CRISRP-mediated CCDC32 knockout cells, AP2 complexes were severely depleted, and CME was strongly inhibited. They further showed *in vitro,* that CCDC32 was recruited to the AAGAB:α:σ2 hemicomplex, where it displaced AAGAB, recruited µ2 and β2 subunits to assemble the mature AP2 complex and then was released. They were unable to detect interactions between a C-terminally tagged CCDC32 and the mature AP2 complex (Wan et al., 2024).

Using a combination of biochemistry, quantitative live cell imaging and ultrastructure electron microscopy, we report that a functional N-terminally tagged CCDC32 interacts with the intact AP2 complex and is recruited to CCPs. siRNA knockdown of CCDC32 inhibits CCP invagination and stabilization, without impacting the levels of AP2 expression. Thus, we demonstrate a second, critical role for CCDC32 in regulating early stages of CME by regulating the stabilization and invagination of CCPs.

## Results

### CCDC32 is recruited to CCPs

We first explored a direct role for CCDC32 in CME by testing if and when CCDC32 is recruited to CCPs. ARPE-HPV cells that stably express mRuby-clathrin light chain a (mRuby-CLCa) and siRNA-resistant, N-terminally GFP tagged full-length CCDC32 (eGFP-CCDC32(FL)) were generated (Fig. S1) and imaged using Total Internal Reflection Fluorescence Microscopy (TIRFM). In addition to diffuse staining across the inner plasma membrane (PM) surface, suggestive of direct membrane binding, we also observed colocalization of eGFP-CCDC32(FL) and mRuby-CLCa (Fig. 1A) at clathrin coated pits.

**Figure 1:**
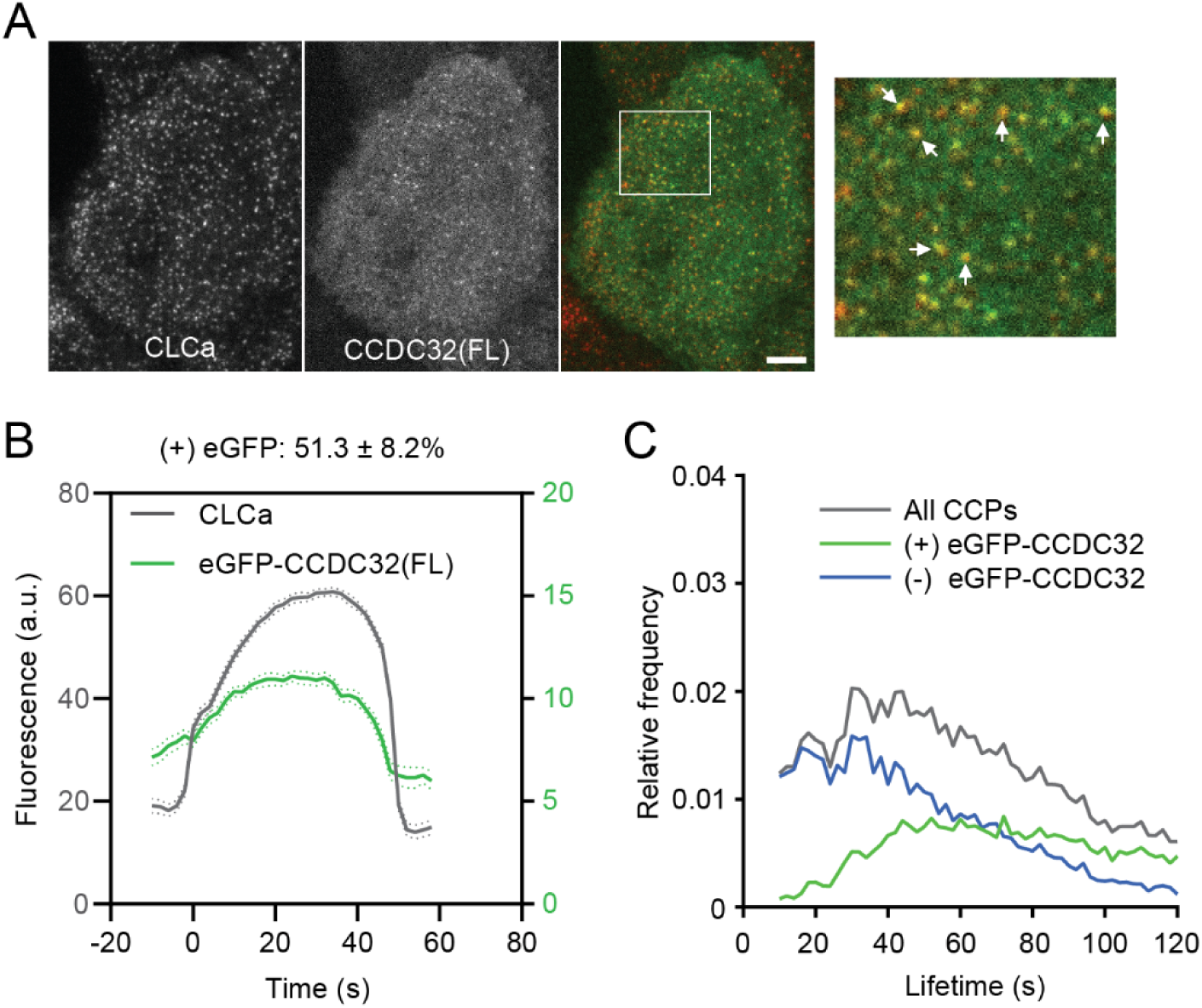
CCDC32 is recruited to clathrin-coated pits. **(A)** Representative TIRFM images of ARPE-HPV cells that stably express mRuby-CLCa and eGFP-CCDC32(FL). This dual channel imaging was conducted without siRNA-mediated knockdown. White arrows point to colocalized CLCa and CCDC32 clusters. White ROI is magnified on the right. Scale bar = 5µm. **(B)** Cohort-averaged fluorescence intensity traces of CCPs (marked with mRuby-CLCa) and CCP-enriched eGFP-CCDC32(FL). 51.3±8.2% of analyzed CCPs showed eGFP-CCDC32(FL) recruitment. Number of tracks analyzed: 23699. **(C)** Lifetime distributions of all CCPs, CCPs with eGFP-CCDC32 recruitment, and CCPs without eGFP-CCDC32 recruitment.

Furthermore, cohort-averaged fluorescence intensity traces were obtained using time-lapse imaging and primary (mRuby-CLCa)/subordinate (eGFP-CCDC32) tracking powered by cmeAnalysis (Aguet et al., 2013; Chen et al., 2020; Jaqaman et al., 2008). As a negative control, ARPE-HPV cells that stably express mRuby-CLCa and eGFP showed neither diffuse PM staining nor eGFP recruitment to CCPs (Fig. S2), whereas more than half of analyzed CCPs were observed to recruit CCDC32. The recruitment curve of CCDC32 follows the assembly of clathrin on CCPs (Fig. 1B). Importantly, compared to CCDC32-positive CCPs, those that fail to recruit eGFP-CCDC32 exhibited significantly shorter, and exponentially decaying lifetimes, previously shown to be characteristic of abortive CCPs (Fig. 1C) (Aguet et al., 2013). Together these data establish that the early recruitment, along with clathrin, of CCDC32 to nascent CCPs regulates their maturation and lifetimes.

### CCDC32 depletion inhibits transferrin receptor (TfnR) uptake and CCP formation

To gain further insight into the function of CCDC32 in CME, we next examined the effects of siRNA-mediated knockdown of CCDC32 on CME and CCP dynamics. In agreement with the previous observation (Wainberg et al., 2021), siRNA-mediated knockdown of CCDC32 in ARPE-HPV cells, which resulted in an ∼60% depletion of CCDC32 (Fig. 2A,B), reduced the cellular uptake efficiency of TfnR (Fig. 2C).

**Figure 2:**
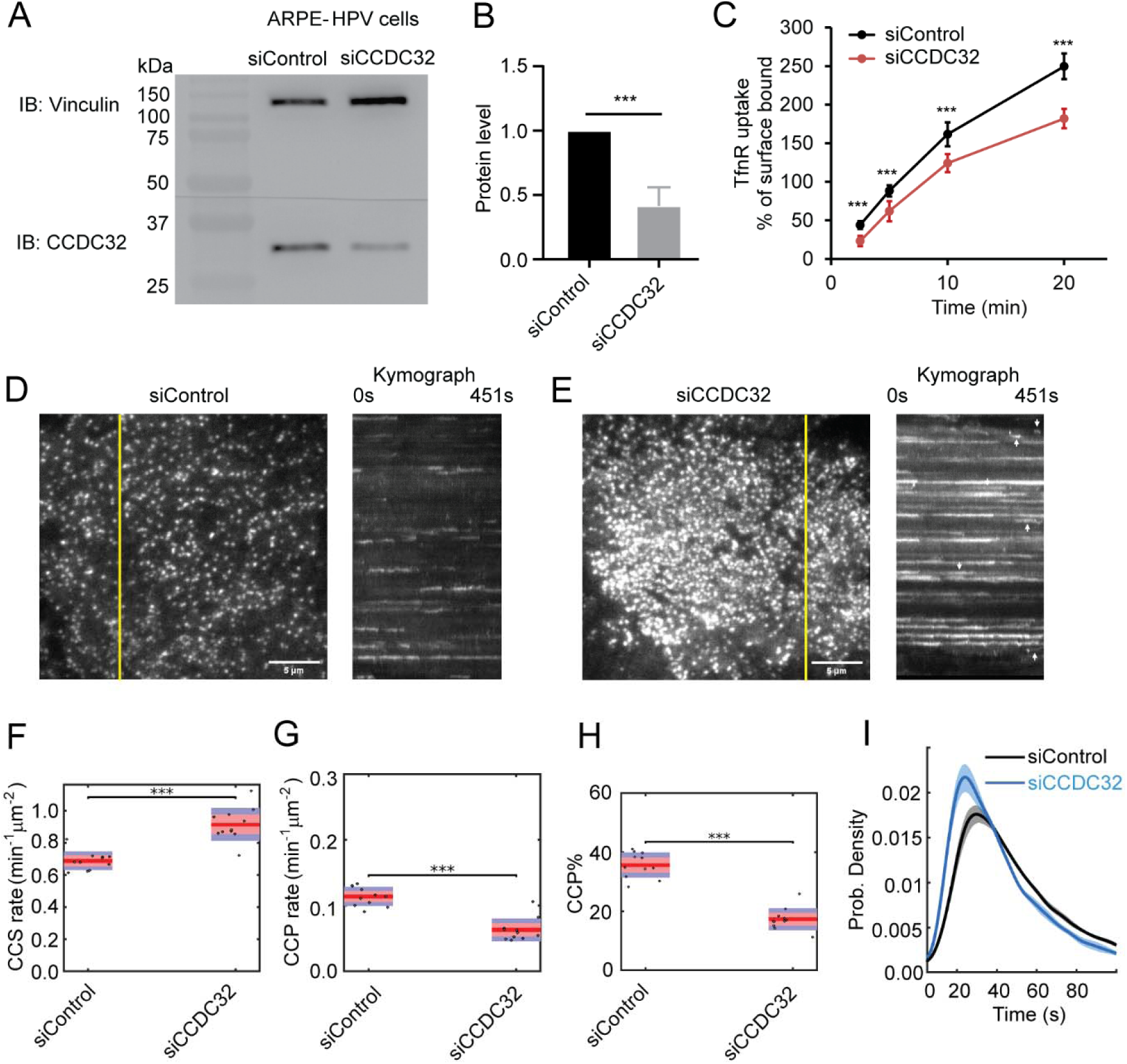
CCDC32 depletion inhibits Transferrin Receptor (TfnR) uptake and CCP maturation. **(A)** Immunoblotting (IB) shows efficient CCDC32 knockdown in ARPE-HPV cells by siRNA treatment. **(B)** Quantified knockdown efficiency (∼60%) of CCDC32 (n=8). **(C)** Measurements of the uptake efficiency of TfnR (n=8). % of surface bound = Internalized/Surface bound*100%. Error bars in (B-C) indicate standard deviations. **(D-E)** Representative single frame images from TIRFM videos (7.5 min/video, 1 frame/s, see videos 1 and 2) and corresponding kymographs from region indicated by yellow lines of ARPE-HPV eGFP-CLCa cells treated with (D) control siRNA or (E) CCDC32 siRNA. Scale bars = 5µm. **(F-H)** Effect of CCDC32 knockdown on the initiation rates of (F) all CCSs and (G) *bona fide* CCPs, as well as (H) the % of *bona fide* CCPs. Each dot represents a video (n=11). Statistical analysis of the data in (F-H)) is the Wilcoxon rank sum test, ***, p ≤ 0.001. **(I)** Lifetime distribution of *bona fide* CCPs. Data presented were obtained from a single experiment (n = 11 videos for each condition) that is representative of 3 independent repeats. Number of dynamic tracks analyzed: 125897 for siControl and 105313 for siCCDC32. Shadowed area indicates 95% confidential interval.

A recent study reported that CCDC32 functions as an essential chaperone for AP2 complex assembly and that CRISPR-mediated knockout of CCDC32 led to a severe decrease in AP2 (Wan et al., 2024). Under our conditions of CCDC32 knockdown, we did not detect any decrease in protein levels of the AP2 complex (Fig. S3A,B), suggesting that the residual CCDC32 was fully capable of fulfilling this catalytic function.

To further define which stages of CME were dependent on CCDC32, we knocked down CCDC32 in ARPE-HPV cells that stably express eGFP-CLCa and employed quantitative live-cell TIRFM to visualize and analyze CCP dynamics (Mettlen and Danuser, 2014). The videos and kymographs of time-lapse imaging showed that CCDC32 depletion resulted in the formation of brighter, static clathrin-coated structures (CCSs) (Fig. 2D,E & Videos 1,2) that dominate the images. These have been seen under other perturbation conditions (Aguet et al., 2013; Chen et al., 2020) and reflect the accumulation of CCPs that are either larger, flatter or both. However, we also noted a subpopulation of dynamic CCPs (arrows, Fig. 2E) that were visually obscured by bright static CCSs, yet represent the majority of total CCSs detected. Indeed, despite the deceptive nature of the images, which we have previously reported (Chen et al., 2020), the percentage of static CCSs (lifetime > 150s), as determined by unbiased quantitative analysis, was only 7.9% (Fig. S4).

To quantify the dynamic behaviors of CCPs, we performed cmeAnalysis (Aguet et al., 2013; Jaqaman et al., 2008; Loerke et al., 2011) and DASC (disassembly asymmetry score classification) (Wang et al., 2020), which together provide a comprehensive and unbiased characterization of CCP intermediates and CME progression when CCDC32 was depleted. DASC, which measures fluctuations of clathrin-assembly/disassembly to accurately distinguish abortive from productive CCPs (Wang et al., 2020), revealed an increased rate of nascent clathrin assembly, reported as the initiation rate of clathrin-coated structures (CCSs) (Fig. 2F), but a reduced rate of *bona fide* CCP initiation (Fig. 2G) and a correspondingly lower percentage of stable, *bona fide* CCPs (CCP%) (Fig. 2H), which are both brighter and longer-lived than abortive CCPs (Wang et al., 2020). Together these data reveal an early defect in the growth and stabilization of nascent clathrin assemblies leading to an increase in the fraction of abortive CCPs. Importantly, the remaining dynamic population of *bona fide* CCPs exhibited shorter lifetimes upon CCDC32 knockdown (Fig 2I), indicating their more rapid maturation.

Together these results demonstrate that CCDC32 is an important endocytic accessory protein involved in CCP stabilization and maturation. Strikingly, despite the profound effects of CCDC32 depletion on CCP dynamics, the efficiency of TfnR uptake was only marginally affected (Fig. 2C). These paradoxical effects are typical for endocytic accessory proteins and indicative of their functional redundancy and/or the induction of compensatory mechanisms, including in this case, the observed increased rate of CCS assembly (Fig. 2E) and the more rapid maturation of *bona fide* CCPs (Fig 2I) (Aguet et al., 2013; Bhave et al., 2020; Wang et al., 2020).

### CCDC32 regulates CCP invagination

During CCP maturation, successful invagination of the clathrin coat and its underlying membrane has been identified as a key step that determines the fate of CCPs (Aguet et al., 2013; Wang et al., 2020). To determine whether CCDC32 regulates CCP invagination, we first used Epifluorescence (Epi)-TIRF microscopy to measure CCP invagination in live cells (Fig. 3A). In this approach, time-lapse Epi and TIRF fluorescence signals were near-simultaneously acquired for ARPE-HPV eGFP-CLCa cells (Fig. 3B) and analyzed using primary (TIRF-channel)/subordinate (Epi-channel) tracking powered by cmeAnalysis (Wang et al., 2020). The resulting Epi and TIRF fluorescence intensity traces of CCPs with similar lifetimes were then aligned and averaged to yield intensity cohorts, which were further log-transformed to give average traces of the invagination depth (Δz) of the CCPs’ center-of-mass (Fig. 3A,C). Here, we chose to present CCPs with lifetimes of 25-35s because they represent the invagination behavior of the most frequent tracks in control cells (Fig. 2H). CCDC32 knockdown strongly inhibited CCP invagination (Fig. 3C), suggesting a key role for CCDC32 in regulating this critical early stage in CME.

**Figure 3:**
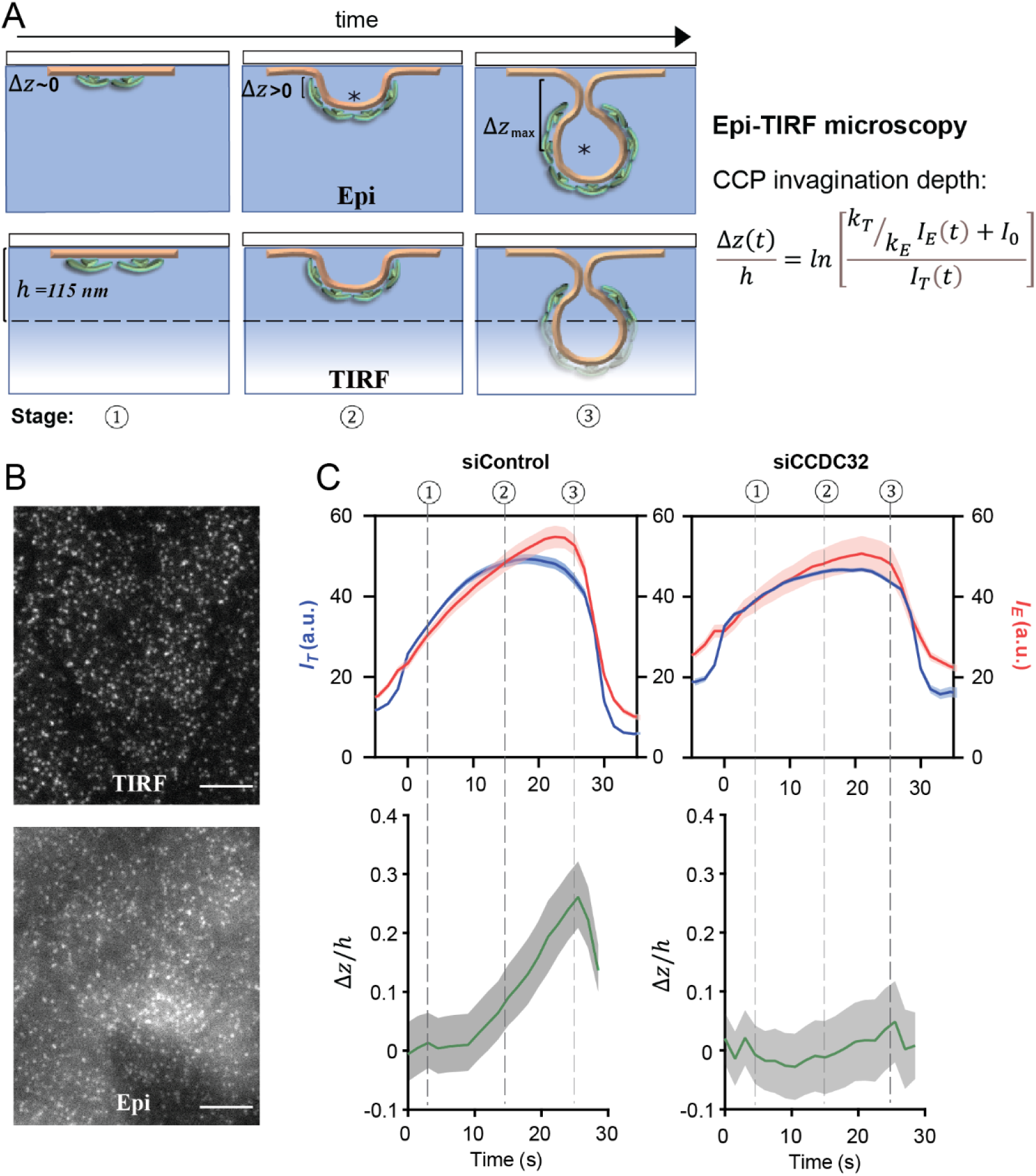
CCDC32 depletion inhibits CCP invagination. **(A)** Scheme of Epi-TIRF microscopy for measuring the invagination of CCPs using primary/subordinate tracking. Δz(t) denotes the invagination depth of CCPs over time. *I_E_* and *I_T_* denotes the cohort-averaged fluorescence intensity from Epi and TIRF channels, respectively. *k_E_* and *k_T_* are the initial growth rate for the Epi and TIRF channel signals, respectively. *I_O_* is an additive correction factor. ‘*’ indicates the mass center of clathrin coat. h=115nm is the evanescent depth of TIRF field, see more detail in ref.(Saffarian and Kirchhausen, 2008; Wang et al., 2020). **(B)** Representative Epi and TIRF microscopy images. Scale bars = 10µm. **(C)** Epi-TIRF microscopy analysis shows that knockdown CCDC32 strongly inhibited CCP invagination. Top: Cohort-averaged CCP fluorescence intensity traces from Epi and TIRF channels; bottom: calculated Δ*z*(*t*)/*h* curves. Data presented were obtained from n = 12 videos for each condition. Number of CCP tracks analyzed to obtain the Δ*z*/*h* curves: 5998 for siControl and 4418 for siCCDC32. Shadowed area indicates 95% confidential interval.

To independently validate that CCDC32 regulates CCP invagination, we next examined the clathrin-coated structures (CCSs) at higher resolution using Platinum Replica Electron Microscopy (PREM) (Svitkina, 2017). Strikingly, CCDC32 knockdown resulted in a substantial increase in the number of flat CCSs (Fig. 4A-C, pseudo-colored blue in panel B, see also Fig. S5A-C) and a corresponding higher occupancy of CCSs on the PM (Fig. 4E; Fig. S5D), consistent with TIRFM and Epi-TIRF microscopy observations. While the flat CCSs we detected in CCDC32 knockdown cells were significantly larger than in control cells (Fig. 4D, mean diameter of 147 nm vs. 127 nm, respectively), they are much smaller than typical long-lived flat clathrin lattices (d≥300 nm) (Grove et al., 2014). Indeed, the surface area of the flat CCSs that accumulate in CCDC32 KD cells (mean ∼1.69 x 10^4^ nm^2^) remains significantly less than the surface area of an average 100 nm diameter CCV (∼3.14 x 10^4^ nm^2^). Thus, we refer to these structures as ‘flat clathrin assemblies’ because they are neither curved ‘pits’ nor large ‘lattices’. Rather, the flat clathrin assemblies represent early, likely defective, intermediates in CCP formation. Importantly, while significantly decreased in both number and size, dome-shaped (green) and spherical (orange) CCSs were still detected, likely corresponding to intermediates within the larger subpopulation of dynamic, *bona fide* CCPs able to maintain TfnR uptake (Fig. 4A-D; Fig. S5A-C). The PREM results provide high resolution structural data supporting a critical role for CCDC32 in regulating CCP invagination.

**Figure 4:**
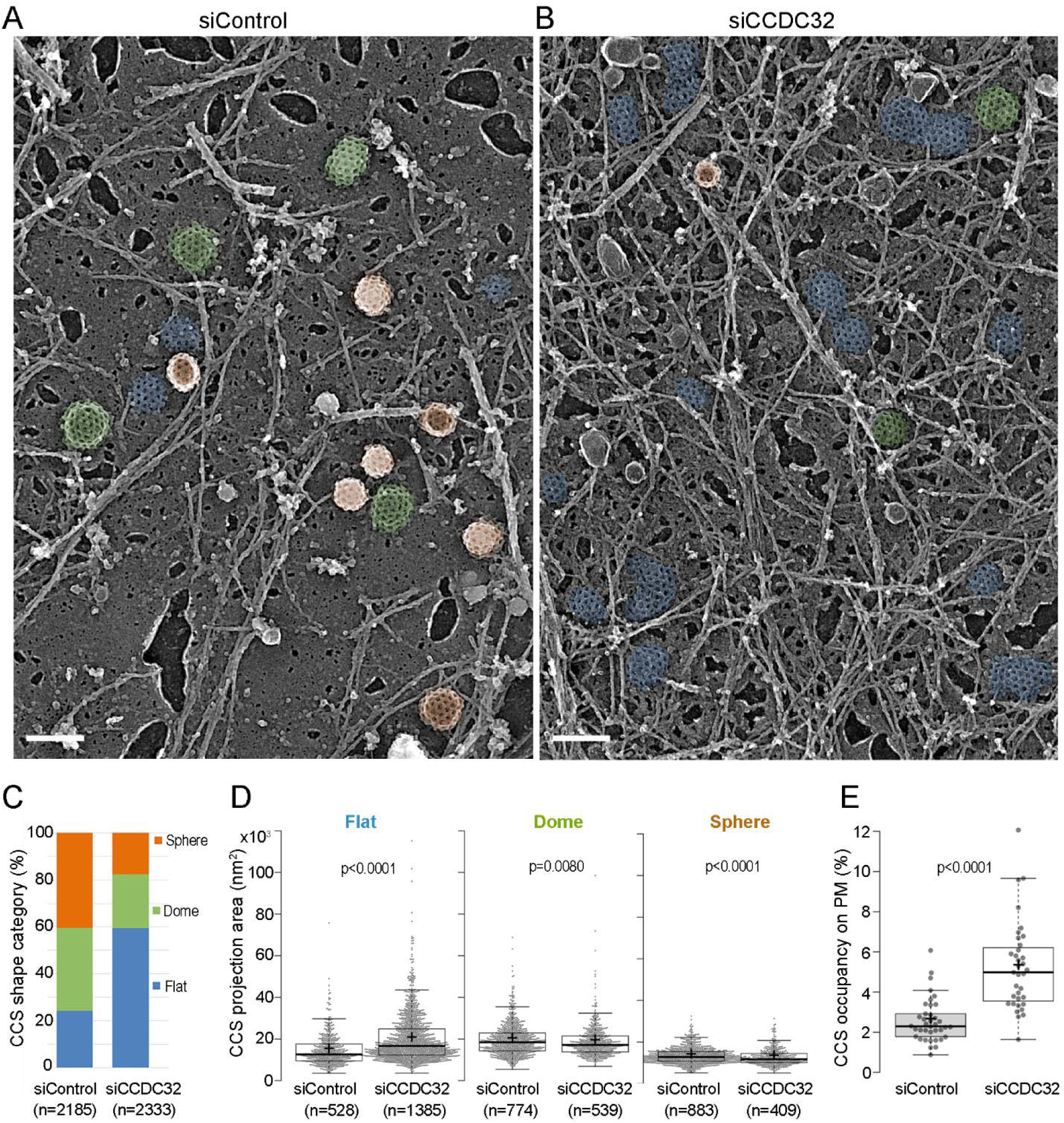
Flat clathrin lattices accumulate in cells depleted of CCDC32. (A-B) Representative PREM images of ARPE19 cells treated with (A) control siRNA or (B) CCDC32 siRNA showing flat (blue), dome-shaped (green) and spherical (orange) CCSs. Scale bars = 200nm. **(C-E)** Quantification of the clathrin-coated structures (CCSs) by (C) shape category, (D) the CCS projection area, and (E) the CCS occupancy on the plasma membrane (PM). Each dot in (E) represents an individual fragment of the PM; the number of cell membrane fragments analyzed is 38 for siControl cells and 35 for siCCDC32 cells from two independent experiments, n = number of CCS. Statistical tests were performed using Mann-Whiney test (D) or unpaired t-test (E). For the Box and whisker plots in (D) and (E), the box extends from the 25th to 75th percentiles, the line in the middle of the box is plotted at the median, and the “+” indicates the mean.

### CCDC32 interacts with the AP2 α-appendage domain

Next, we explored the mechanism of CCDC32 recruitment to CCPs. The adaptor AP2, which is essential for CCP and CCV formation, is a heterotetramer consisting of α, β2, μ2, and σ2 subunits (Fig. 5A). A previous study had reported interactions between overexpressed σ2-mCherry and CCDC32-GFP (Wainberg et al., 2021), but had not demonstrated interactions with the native AP2 complex. A more recent study reported that C-terminally tagged CCDC32 interacts with the α:σ2 hemicomplex, but does not interact with the assembled AP2 heterotetramer (Wan et al., 2024). To explore this apparent discrepancy, we conducted co-immunoprecipitation (co-IP) experiments from cell lysates of ARPE-HPV cells that stably express a fully functional eGFP-tagged α-subunit of AP2 (Mino et al., 2020) (AP2-α-eGFP, Fig. 5B). Mass Spectrometry analysis revealed, as expected, the co-IP of all the subunits of the AP2 complex, as well as well-known AP2 binding proteins (i.e. EPS15, NECAP2, AAK1 and AAGAB) (Mettlen et al., 2018). Notably, CCDC32 was also efficiently co-immunoprecipitated with intact AP2 (Fig. 5C; Table S1). However, this experiment does not definitively rule out the possibility that CCDC32 only interacts with a small population of immature α:σ2 hemicomplexes that might be present in the lysate. Therefore, we performed co-IP experiments in cell lysates from ARPE-HPV cells stably expressing eGFP-CCDC32(FL) (Fig. 5D). Western blotting showed that the α subunit (Fig. 5E-F), together with each of AP2 complex subunits (Fig. S3C) were efficiently co-precipitated with eGFP-CCDC32(FL), but not by GFP (Fig. 5D-F). Thus, we conclude that CCDC32 interacts with the mature AP2 complex. The discrepancy with previous findings may be due to the location of the eGFP tag (see discussion).

**Figure 5:**
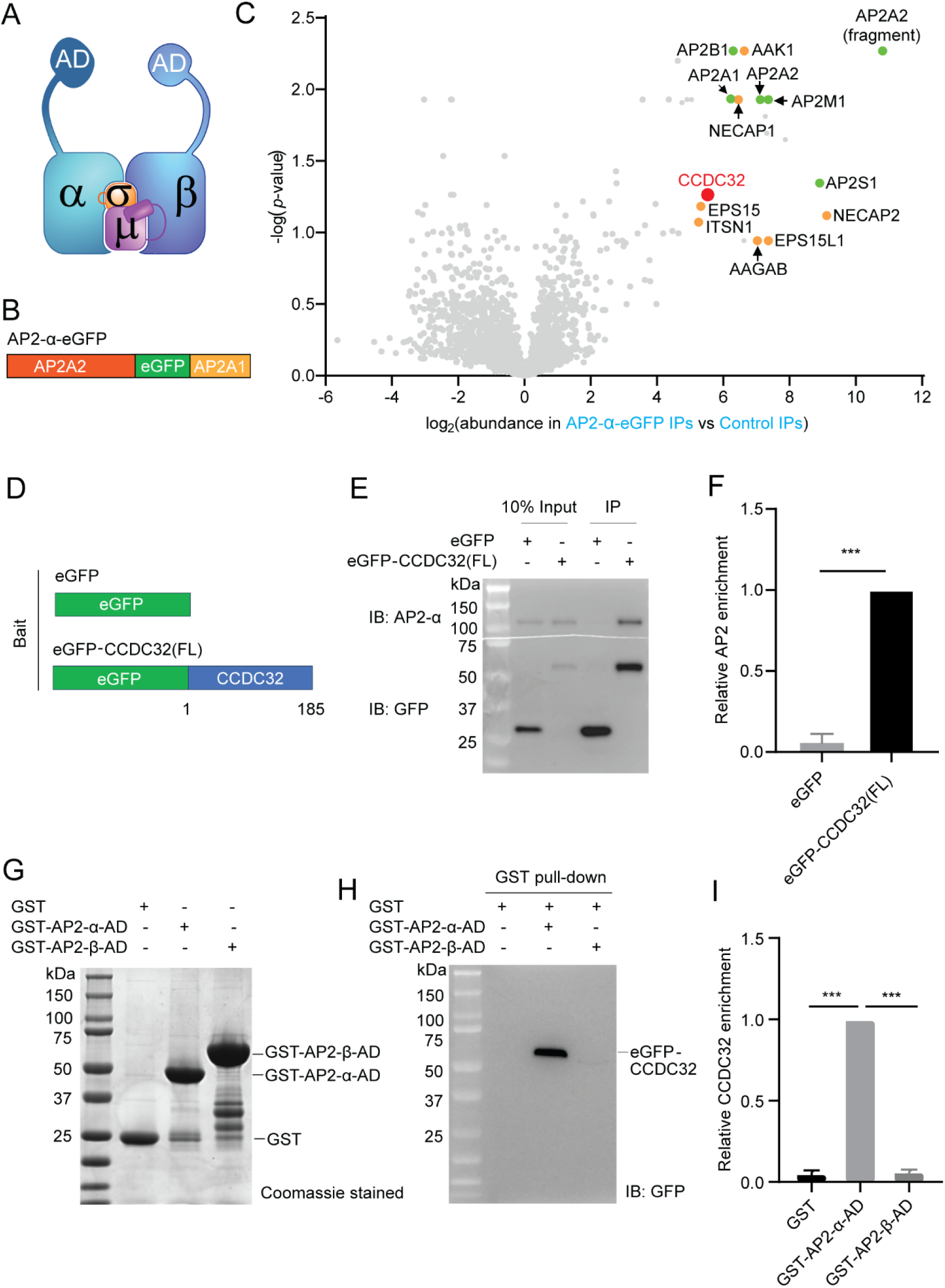
CCDC32 interacts with AP2 through the α appendage domain. (A) AP2 structure. AD: appendage domain. **(B)** Domain structure of a fully-functional α-subunit bearing an internal eGFP tag inserted in the unstructured hinge region. **(C)** Volcano plot after mass spectrometry analysis of the immunoprecipitation (IP) of eGFP from naive ARPE-HPV cells (control IPs) or ARPE-HPV cells that stably express AP2-α-eGFP (AP2-α-eGFP IPs) using anti-GFP beads. Green: AP2 subunits; orange: known AP2 interactors. Source data for the volcano plot is available as Supplementary Table 2. **(D-F)** IP of eGFP from ARPE-HPV cells that stably express eGFP or eGFP-CCDC32(FL) using anti-GFP magnetic beads. (D) The domain structures of eGFP and eGFP-CCDC32(FL). (E) Representative immunoblotting result of n=3 IP samples. (F) Relative AP2 enrichments quantified from immunoblotting results. **(G-I)** GST pull-down assays. (G) Quantification of immunoblots of the relative enrichment of CCDC32. (H) Coomassie blue stained SDS-page gel of purified GST, GST-AP2-α-AD, and GST-AP2-β-AD. (I) Representative western blot result of n=3 GST pull-down assay from ARPE-HPV eGFP-CCDC32(FL) cell lysate using purified GST, GST-AP2-α-AD, or GST-AP2-β-AD. The amount of bait GST-proteins is shown in (D), and the pulled down eGFP-CCDC32 was detected by immunoblotting (IB) of GFP. Error bars in (F) and (I) are SD (n=3).Two-tailed student’s t-test: ***, p ≤ 0.001.

Most endocytic accessory proteins interact with AP2 via the appendage domains of the α and β2 subunits (Praefcke et al., 2004; Schmid et al., 2006) (Fig. 5A, α-AD and β-AD). Therefore, we expressed and purified GST-tagged AP2 α-AD and β-AD (Fig. 5G) and used GST pull-down assays to test which might interact with CCDC32. GST-AP2-α-AD, but not GST-AP2-β-AD, was capable of pulling down CCDC32 from ARPE-HPV eGFP-CCDC32(FL) cell lysate (Fig. 5H,I). Thus, CCDC32 can interact with AP2 via the α-AD.

### Identification of a CCDC32 binding site for AP2 interactions

We next identified a region on CCDC32 responsible for AP2 binding. Structural predictions of CCDC32 made by AlphaFold 3.0 show it to be a mainly unstructured protein with several isolated α-helices (Fig. S6A,B), which likely led to its misnomer as a coiled-coil domain containing protein. Indeed, even when modelled as a dimer or trimer, we found no evidence of coiled-coil interactions between the α-helices (data not shown). Moreover, endogenous CCDC32 did not co-immuno-precipitate with eGFP-CCDC32 from ARPE-HPV eGFP-CCDC32(FL) cell lysate (Fig. S6C), indicating that the protein is a monomer *in vivo*. Nonetheless, we focused first on a strongly predicted α-helix encoded by aa78-98 and located in the middle of CCDC32 and engineered a CCDC32 construct lacking this α-helix (Δ78-98, Fig. 6A; Fig. S6A-B). The corresponding siRNA-resistant CCDC32(Δ78-98) mutant was stably expressed in ARPE-HPV cells at comparable expression levels to eGFP-CCDC32(FL) (Fig. S1A). As previously showed (Fig. S3C) all four subunits of AP2 efficiently co-IP with full length CCDC32 from cell lysates (Fig. 6B,C). Interestingly, the co-IPs of both α and σ2 subunits with CCDC32(Δ78-98) were greatly reduced (Fig. 6B,C); whereas, unexpectedly the ability of CCDC32(Δ78-98) to interact with β2:µ2 was unaffected. These data are partially consistent with the results of Wan et al.(Wan et al., 2024), who reported interactions with the α:σ2 hemicomplex and with µ2, but not with β2. Given the efficiency and selectivity of our co-IP, our data also suggests that the mature AP2 complex exists in a dynamic equilibrium between α:σ2 and β2:µ2 hemicomplexes. . Indeed, it has previously been reported that β2:µ2 hemicomplexes can partially support synaptic vesicle recycling in C. elegans bearing null mutations in the α-subunit of AP2(Gu et al., 2013). These findings demonstrate that aa78-98 are essential for CCDC32 interactions with both mature AP2 complexes and the α:σ2 hemicomplex.

**Figure 6:**
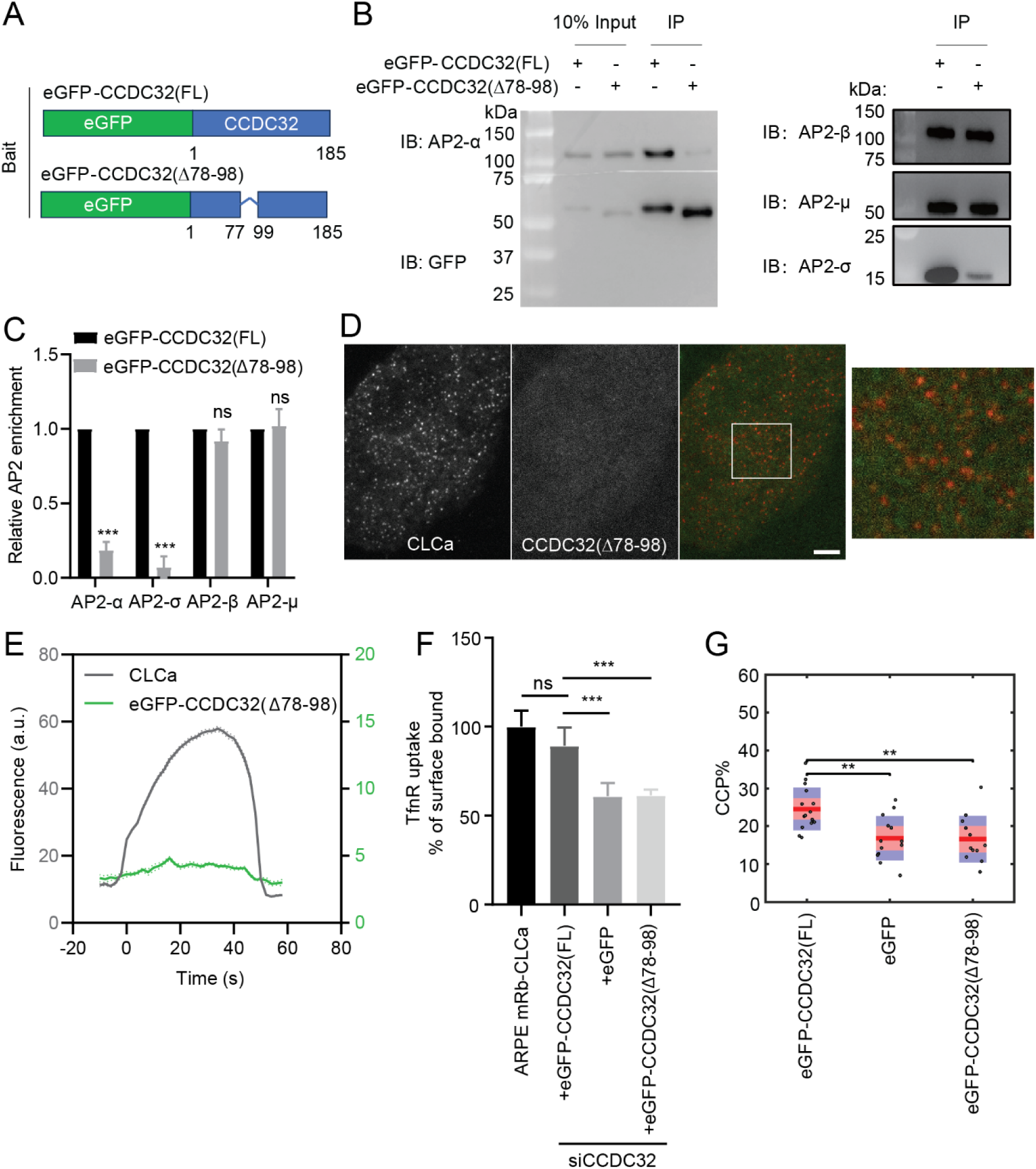
A central α-helix in CCDC32 mediates CCDC32-AP2 interactions. (A-C) IP of eGFP from ARPE-HPV cells that stably express eGFP-CCDC32(FL) or eGFP-CCDC32(Δ78-98) using anti-GFP beads. (A) The domain structures of eGFP-CCDC32(FL) and eGFP-CCDC32(Δ78-98). (B) Representative Immunoblotting result of n=3 IP samples. (C) Relative AP2 enrichments that were quantified from (B). Error bars indicate standard deviations. **(D)** Representative TIRFM images of ARPE-HPV cells that stably express mRuby-CLCa and eGFP-CCDC32(Δ78-98). White ROI is magnified on the right. Scale bar = 5µm. **(E)** Cohort-averaged fluorescence intensity traces of CCPs and CCP-enriched eGFP-CCDC32(Δ78-98). Number of tracks analyzed: 37892. **(F-G)** (F) TfnR uptake efficiency and (G) CCP% of ARPE mRb-CLCa cells that stably express eGFP-CCDC32(WT), eGFP, or eGFP-CCDC32(Δ78-98), and with siRNA-mediated knockdown of endogenous CCDC32. Each dot in (G) represents a movie. Statistical analysis of the data in (F) is two-tailed student’s t-test: ns, not significant; ***, p ≤ 0.001. Statistical analysis of the data in (G) is the Wilcoxon rank sum test, **, p ≤ 0.01.

Disrupting AP2 interactions severely impaired the recruitment of CCDC32(Δ78-98) to CCPs (Fig. 6D,E), indicating that CCDC32 is likely recruited to CCPs through its interactions with AP2. Notably, unlike CCDC32(FL), CCDC32(Δ78-98) was unable to rescue either the uptake of the CME cargo, TfnR (Fig. 6F) or the stabilization of CCPs (Fig. 6G). Together, these results establish that CCDC32, a previously uncharacterized endocytic accessory protein, plays a critical role in regulating CCP stabilization and invagination.

### Disease-causing nonsense mutation in CCDC32 loses AP2 interaction capacity and inhibits CME

Loss-of-function nonsense mutations in CCDC32 have been reported to result in CFNDS (Abdalla et al., 2022; Harel et al., 2020); however, the disease-causing mechanism remains unknown. The identified frameshift mutations result in premature termination and truncation of the protein at residue 10, 55 or 81 (Harel et al., 2020). Based on our findings, we hypothesize that these C-terminally truncated mutants, all of which lack the α-helix encoded by residues 78-98, will be defective in AP2 binding and unable to function in CME. To test this hypothesis, we generated an siRNA-resistant eGFP-CCDC32(1-54) construct (Fig. 7A) based on the clinical report (Harel et al., 2020), and stably expressed this truncated CCDC32 mutant in ARPE-HPV cells (Fig. S1A).

**Figure 7:**
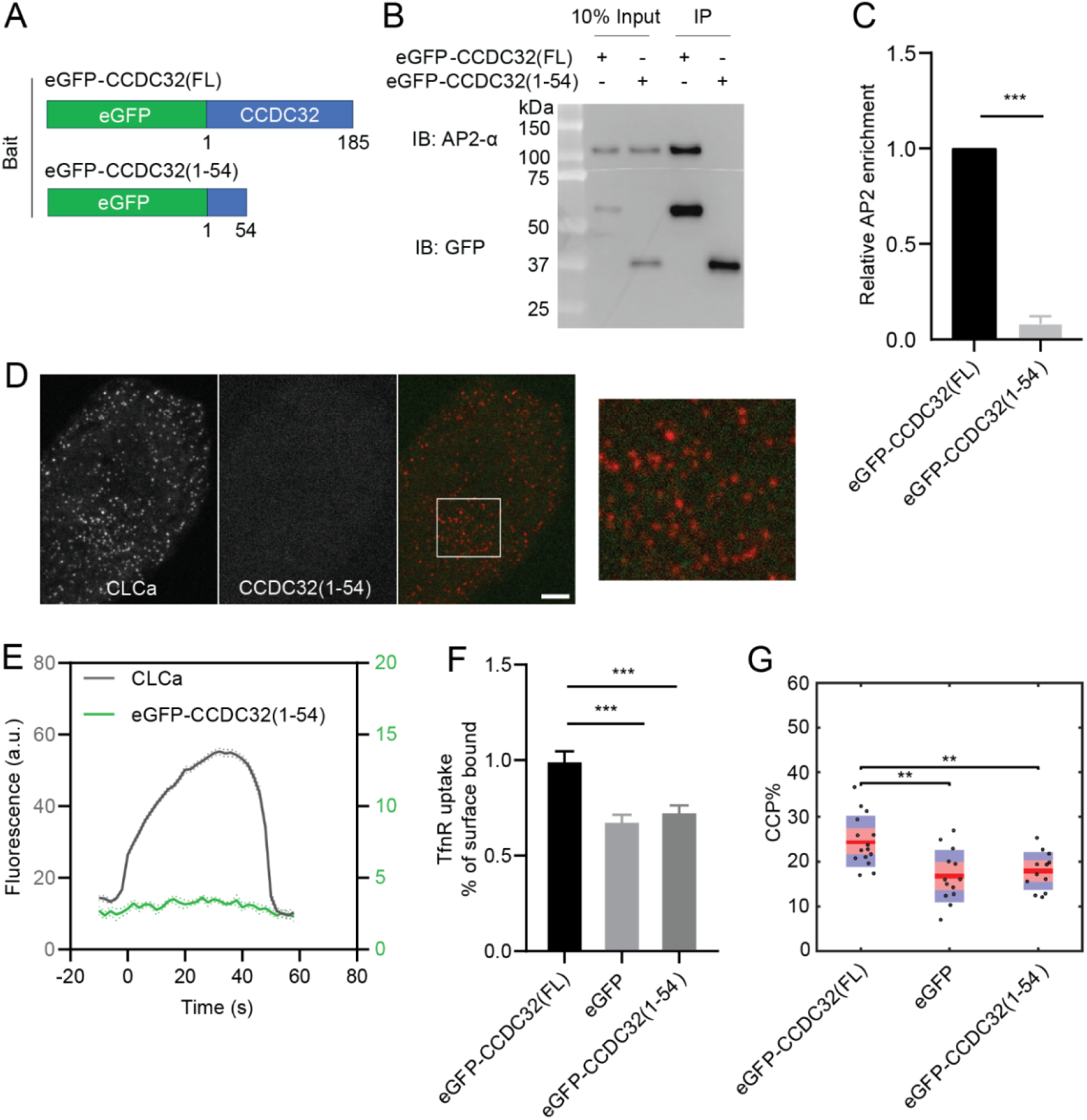
Disease-causing nonsense mutation in CCDC32 loses AP2 interaction capacity and inhibits CME. (A-C) IP of eGFP from ARPE-HPV cells that stably express eGFP-CCDC32(FL) or eGFP-CCDC32(1-54) using anti-GFP magnetic beads. (A) The domain structures of eGFP-CCDC32(FL) and eGFP-CCDC32(1-54). (B) Representative immunoblotting result of n=3 IP samples. (C) Relative AP2 enrichments quantified from immunoblotting results. **(D)** Representative TIRFM images of ARPE-HPV cells that stably express mRuby-CLCa and eGFP-CCDC32(1-54). White ROI is magnified on the right. Scale bar = 5µm. **(E)** Cohort-averaged fluorescence intensity traces of CCPs and CCP-enriched eGFP-CCDC32(1-54). Number of tracks analyzed: 28658. **(F-G)** (F) TfnR uptake efficiency and (G) CCP% of ARPE-HPV cells that stably express eGFP-CCDC32(FL), eGFP, or eGFP-CCDC32(1-54), and with siRNA-mediated knockdown of endogenous CCDC32. Each dot in (G) represents a movie. Statistical analysis of the data in (F) is two-tailed student’s t-test: ***, p ≤ 0.001. Statistical analysis of the data in (G) is the Wilcoxon rank sum test, **, p ≤ 0.01.

As predicted, AP2 did not co-IP with CCDC32(1-54) from the cell lysate (Fig. 7B,C), confirming that the disease-causing mutation in CCDC32 abolishes its interactions with AP2. Correspondingly, we could not detect CCDC32(1-54) recruitment to CCPs in dual channel TIRFM imaging (Fig. 7D,E), consistent with CCP recruitment of CCDC32 being dependent on CCDC32-AP2 interactions. Interestingly, we also noticed that the level of diffuse plasma membrane binding detected by TIRFM decreased to a greater extent than that seen for CCDC32(Δ78-98), now corresponding to background, eGFP only levels (see Fig. S7 for direct comparison). These data suggest that the C-terminus of CCDC32 is required for binding to the inner surface of the plasma membrane. Finally, we observed that expression of eGFP-CCDC32(1-54), after siRNA-mediated knockdown of endogenous CCDC32, was unable to rescue TfnR uptake efficiency (Fig. 7F) or CCP stabilization (Fig. 7G). These findings show that this loss-of-function nonsense mutation in CCDC32 abolishes its interactions with AP2 and inhibits CME likely contributing to the development of CFNDS.

## Discussion

CCDC32 was identified through genetic studies, originally in a yeast 2-hybrid screen for proteins interacting with annexin A2 (Li et al., 2011) and subsequently via whole exome sequencing to identify mutations associated with cranio-facio-neurodevelopmental syndrome (CFNDS) (Abdalla et al., 2022; Harel et al., 2020), neither of which provided insight into its cellular function. Subsequent bioinformatic analysis of co-essential modules linked CCDC32 with the AP2 adaptor complex, provided evidence for its interaction with AP2, reported colocalization of CCDC32 with AP2 at CCPs, and demonstrated a role in CME (Wainberg et al., 2021). A more recent report has suggested that CCDC32 functions as a chaperone, essential for the assembly of mature AP2 heterotetrametric complexes (Wan et al., 2024), but that CCDC32 neither binds to the mature AP2 complex nor colocalizes with CCPs.

Here we show that CCDC32 binds to intact AP2 complexes and that this interaction is required for its recruitment to CCPs. Our data demonstrate that CCDC32 plays a critical role at early stages of CME, dependent on its recruitment to CCPs. Depletion of CCDC32 results in a pronounced defect in CCP invagination, a decrease in the rate of formation and percentage of stabilized nascent *bona fide* CCPs and an accumulation of flat clathrin assemblies, unstable intermediates in CCP formation (Fig. 8). Despite these profound alterations in CCP dynamics, CME itself, as measured by TfnR internalization efficiency, is only partially inhibited. We speculate that this mild endocytic defect reflects the plasticity and resilience of CME. Indeed, we detect two potential compensatory mechanisms that occur upon depletion of CCDC32, namely an increase in the rate of CCS assembly and a decrease in the lifetimes of dynamic *bona fide* CCPs (i.e. an increased rate of CCP maturation) (Fig. 8).

**Figure 8:**
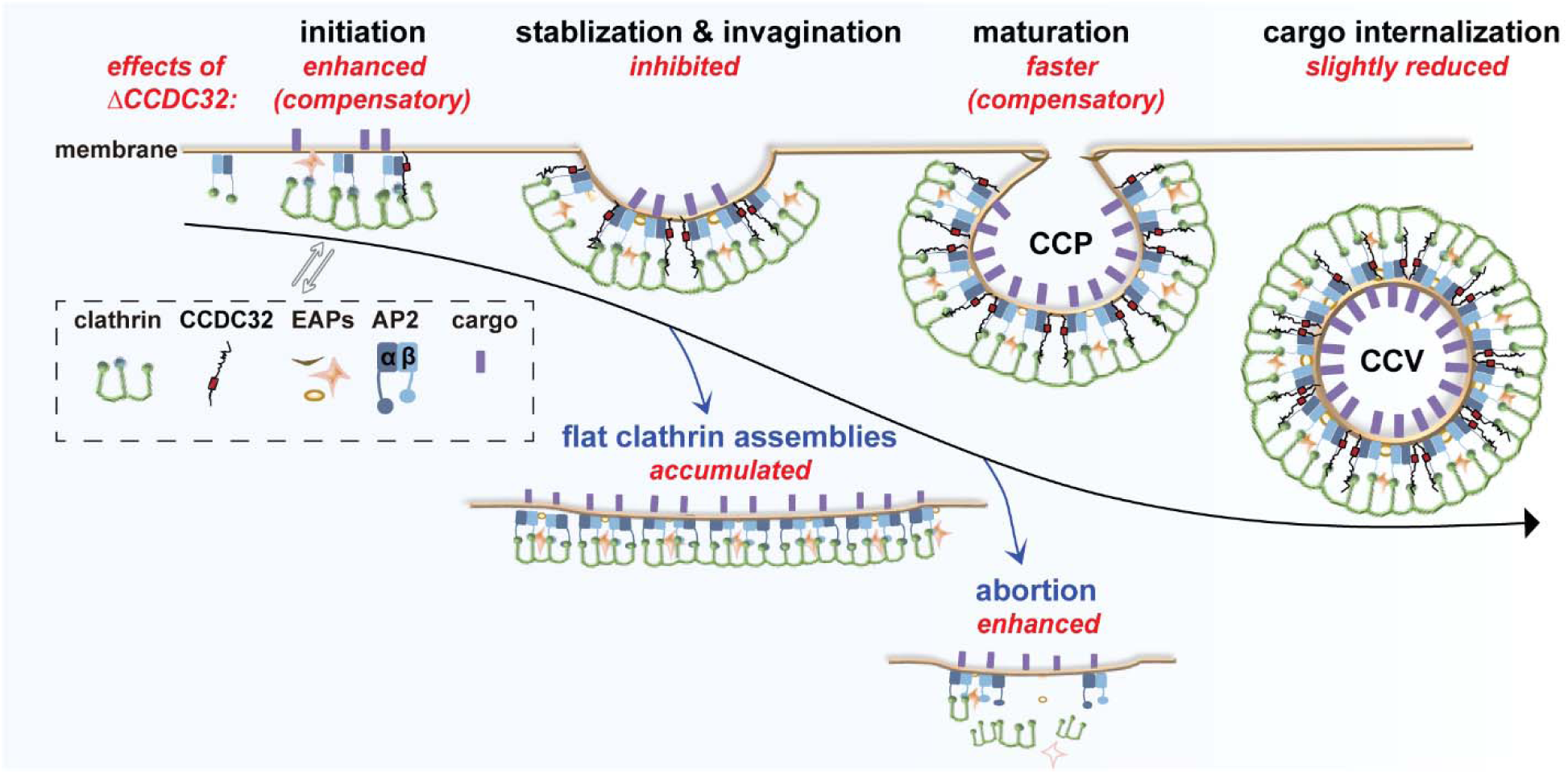
Cartoon illustration of CCDC32-AP2 interactions that regulate CME. Interactions between residues 78-98, corresponding to a central α-helix of CCDC32, and AP2 are essential for recruitment of CCDC32 to CCPs. Depleting CCDC32 (ΔCCDC32) inhibits CCP stabilization and invagination, resulting in enhanced CCS abortion and accumulated flat clathrin assembly intermediates. As a compensation effect, CCS initiation is enhanced and CCP maturation is faster, resulting in only slightly reduced cargo internalization. EAPs: endocytic accessory proteins.

While individual depletion of other endocytic accessory proteins has been reported to result in one or more of these phenotypes, to our knowledge, this combination of phenotypes has not been detected. Instead, either CCP assembly is reduced (Henne et al., 2010; Umasankar et al., 2014) or CCPs accumulate at later stages of invagination (McMahon and Boucrot, 2011; Messa et al., 2014). This combination of phenotypes has, however, been observed previously in cells expressing a truncated mutant of AP2 lacking the α-AD (Aguet et al., 2013). These cells also exhibited: i) near normal Tfn uptake, ii) an accumulation of flat clathrin assemblies, which are rapidly turned over, iii) an increase in the rate of nascent CCS initiation, iv) impaired stabilization of dynamic CCPs and, v) a decrease in productive CCP lifetimes. We speculate that loss of CCDC32 recruitment could have accounted for these unique phenotypes.

Although the mechanism of CCP invagination has been an important topic for decades, which endocytic accessory proteins (if any) regulate CCP invagination and (if so) how, has remained unclear. The depletion of CCDC32 in cells strongly inhibited the formation of dome-shaped and spherical CCSs, and instead resulted in the accumulation of flat clathrin assemblies. As the area of these clathrin lattices was insufficient to form a complete CCV, they likely represent defective CCP assembly intermediates that are rapidly turned over as abortive CCPs. This experimentally observed requirement of CCDC32 in CCP invagination is consistent with previous findings that clathrin assembly alone was not sufficient to induce CCP invagination in cells (Aguet et al., 2013), potentially due to the reported flexibility of clathrin coats (Ferguson et al., 2008; Tagiltsev et al., 2021). The mechanism by which CCDC32 promotes CCP invagination remains to be determined, but other domains of the protein may bind to and stabilize the clathrin coat, or the predicted largely disordered structure of CCDC32 (Fig. S6) may generate membrane curvature by molecular crowding (Busch et al., 2015).

We localized the AP2 binding site on CCDC32, which is essential for CCDC32-AP2 interactions *in vivo*, to a short, predicted alpha-helical region (aa78-98). Interestingly, this sequence, LASLEKKLRRIKGLNQEVTSKD, does not encode any of the known AP2 α-AD binding motifs (e.g. DxF/W, FxDxF, WxxF/W, FxxFxxL) commonly shared amongst other AP2 interacting partners (Olesen et al., 2008). However, we identified two canonical α-AD binding motifs in CCDC32 (^17^DLW^19^ and ^39^FSDSF^43^), both of which can be docked on the AP2 appendage domain with high confidence using AlphaFold 3.0 (Fig. S6D,E). While these binding motifs may have been sufficient for *in vitro* interactions with α-AD, they were not sufficient for AP2 binding *in vivo*. Unexpectedly, AlphaFold 3.0 modeling predicts, with high confidence, that the α-helical region we identified as essential for AP2 binding *in vivo* docks to an alpha helix formed by aa418-436 of the α-subunit (Fig. S6D,F), which is not encoded in our α-AD construct. This site is located in the α-trunk and buried with the AP2 core domain (Collins et al., 2002). However, it becomes accessible when the C-terminal domain of µ2 adopts the open conformation triggered by cargo binding (Jackson et al., 2010). Thus, we speculate that CCDC32 may preferentially interact with membrane-bound AP2. Interestingly, while co-IP with CCDC32(Δ78-99) no longer pulled down the intact AP2 complex or the α:σ2 hemicomplex, it remained capable of pulling down the β2:µ2 hemicomplex. This result has a number of interesting implications: first that the extended full length CCDC32 has multiple points of interaction with AP2 and its subunits, that will require further studies to structurally and functionally define; and second, that the AP2 complex is less stable than previously assumed (i.e. the 2 hemicomplexes must be in flux). Indeed, it has been shown that β2:µ2 hemicomplexes can partially support synaptic vesicle recycling in C. elegans bearing null mutations of the α-subunit (Gu et al., 2013).Further structural studies will be needed to identify the full extent of CCDC32 interactions with the AP2 complex.

While this paper was under review, it was reported that CCDC32 functions together with AAGAB, as an essential chaperone for AP2 assembly (Wan et al., 2024). While there are discrepancies, our results are not incompatible with their findings. Thus, it is possible, even likely, that the ∼40% residual CCDC32 present after siRNA knockdown may be sufficient to fulfill its catalytic chaperone activity in facilitating AP2 complex assembly, but not its presumed structural role in regulating early stages of CME. The inability of Wan et al. to detect CCDC32 binding to mature AP2 complexes or β2:µ2 hemicomplexes or its recruitment to CCPs may reflect a perturbed function of the C-terminally tagged construct (Wan et al., 2024). Indeed, their images show the formation of large (>100 nm), cytoplasmic puncta of CCDC32-GFP, which might be reflective of protein aggregation. Moreover, our data show a role for the C-terminus in diffuse plasma membrane binding. Finally, we find that the Δ78-98 mutant retains its ability to bind the β2:µ2 hemicomplex, but loses its ability to bind α:σ2 hemicomplex. Thus, the most parsimonious conclusion is that CCDC32 is multifunctional, acting catalytically to facilitate AP2 assembly and, after recruitment to CCPs, to regulate early stages of CME. Further studies are needed to fully elucidate the mechanisms governing each of these functions.

Our results provide an explanation for how the clinically observed loss-of-function mutations in CCDC32 result in the development of CFNDS. As we have shown, the disease-associated CCDC32 loss-of-function mutants are not capable of interacting with intact AP2, thus losing their AP2 regulation capacity. Our results suggest that the inability to bind mature AP2 and hence to be recruited to nascent CCSs inhibits critical early stages of CME, and contributes to the development of CFNDS. While we cannot rule of nonsense-mediated decay and resulting loss of expression of the truncated human mutations (Lykke-Andersen and Jensen, 2015), it is interesting that the clinical features of CCDC32 loss-of-function are similar to those resulting from AP2 loss-of-function mutations (Arrigo and Lin, 2021; Gorvin et al., 2017; Helbig et al., 2019; Jung et al., 2015; Li et al., 2018; Li et al., 2010), supporting that CCDC32 functions through AP2. However, we suggest that the disease-associated CFNDS mutants are hypomorphic, as the complete loss of AP2 complexes, which would result from a complete loss of the CCDC32 protein (Wan et al., 2024), has been shown to be embryonic lethal in Drosophila (González-Gaitán and Jäckle, 1997) and zebrafish (Umasankar et al., 2012).

In summary, our study identified CCDC32 as an important endocytic accessory protein that regulates CCP stabilization and invagination via its interactions with AP2. Future work needs to address the multifunctional interactions of CCDC32 with AP2 and to define exactly how CCDC32 enhances curvature generation and coat stabilization.

## Materials and Methods

### Plasmids

The eGFP-CCDC32(FL, human) cDNA in a pEGFP-C1 vector was purchased from Addgene (#110505) and then mutated to be siRNA resistant, which retained amino acid sequence (#98-102) while modifying the nucleotide sequence. Next, aa78-98 was deleted from siRNA resistant eGFP-CCDC32(FL) to generate eGFP-CCDC32(Δ78-98). Finally, eGFP, eGFP-CCDC32(1-54), eGFP-CCDC32(Δ78-98), and eGFP-CCDC32(FL) were separately cloned into a pLVx-CMV100 vector (Dean et al., 2016) using NEBuilder® HiFi DNA Assembly Master Mix (Catalog #E2621). A full list of primers used for mutagenesis and cloning is available in Table S2. Note that our disease mimic construct CCDC32(1-54) does not contain a 9 aa peptide (VRGSCLRFQ) in the N-terminus and an extra 12 aa in the C-terminus when CFNDS patient mutation was described (p.(Glu64Glyfs∗12)) in ref. (Harel et al., 2020).

In addition, mRuby-CLCa in a pLVx-IRES-puro vector was generated in our previous study(Srinivasan et al., 2018). AP2-α-AD (aa701-938, mouse) in a pGEX-2T-1 vector and AP2-β-AD (aa592-937, rat) in a pGEX-4T-1 vector were kind gifts of the late Linton Traub (University of Pittsburgh, PA).

### Cell culture, lentivirus infection, siRNA transfection and rescue

ARPE-19 and ARPE19-HPV16 (herein called ARPE-HPV) cells were obtained from ATCC and cultured in DMEM/F12 (Gibco, Catalog #8122502) with 10% FBS. HEK293T cells were obtained from ATCC and cultured in DMEM (Gibco, Catalog #8122070) with 10% FBS. ARPE-HPV cells that stably express eGFP-CLCa were generated in our previous study (Chen et al., 2020). ARPE-HPV cells that stably express a fully functional AP2-α-eGFP, in which eGFP is inserted into the flexible linker of AP2, at aa649, were generated in our previous study (Mino et al., 2020) and this AP2-α-eGFP construct has been shown to be fully functional.

Lentiviruses encoding mRuby-CLCa were produced in HEK293T packaging cells following standard transfection protocols (Kutner et al., 2009) and harvested for subsequent infections to ARPE-HPV cells to generate ARPE-HPV mRuby-CLCa cells. Lentiviruses encoding eGFP and siRNA-resistant, eGFP-CCDC32(1-54), eGFP-CCDC32(Δ78-98), and eGFP-CCDC32(FL) were produced in HEK293T packaging cells and harvested for subsequent infections to ARPE-HPV mRuby-CLCa cells. Infected cells were FACS sorted for homogenous mRuby and eGFP signals after 3 days and passaged for 2 weeks before experiments.

For siRNA-mediated knockdown of CCDC32, ARPE-HPV cells stably expressing eGFP-CLCa were seeded on 6-well plates (250,000 cells/well) and transfected with 2 rounds of siCCDC32 (Silencer Select Pre-designed siRNA ID#: s228444, targets aa98-102 sequence) through 3 days. Cells treated with siControl (Silencer Select Negative Control #1 siRNA, cat#:4390843) were used as negative control. Transfections of siRNA were mediated with Opti-MEM and Lipofectamine RNAi-MAX (Invitrogen) as detailed in ref. (Chen et al., 2020).

For CCDC32 rescue experiments, ARPE-HPV cells stabily expressing eGFP or siRNA-resistant eGFP-CCDC32(FL) eGFP, eGFP-CCDC32(Δ78-98 or eGFP-CCDC32(1-54) were transfected with siCCDC32 or siControl as described above. Anti-GFP Monoclonal Antibody (Proteintech, # 66002-1-Ig), Anti-Vinculin Polyclonal Antibody (Proteintech, #26520-1-AP), and anti-C15orf57 Polyclonal antibody (Invitrogen, #PA5-98982) were used in Western Blotting to confirm protein expression level and knockdown efficiency.

### Co-immunoprecipitation (co-IP)

ARPE-HPV cells with stable expression of AP2-α-eGFP, eGFP, eGFP-CCDC32(1-54), eGFP-CCDC32(Δ78-98) or eGFP-CCDC32(FL) were cultured in a 15-cm dish. When confluence reached ∼90%, cells were washed 3x with ice-cold PBS and detached with cell scratcher. Next, cells were spun down at 500 g, 4°C for 3 min, and then resuspended in 1ml ice-cold lysis buffer (50 mM Tris, pH 7.5, 150 mM NaCl, 1mM EDTA, 0.5% Triton X-100, 1× protease inhibitor, 1mM PMSF). After 30 min rotation in a cold room and occasional vortexing, the lysed cells were spun at 500 g, 4°C, for 3 min to remove the nuclei. Subsequently, 50µl anti-GFP magnetic beads (Biolinkedin, #L1016) were added to cell lysate that contained 0.5mg proteins (determined by BCA assay). The reaction was allowed to proceed by rotation at 4°C for 2h and then precipitated with DynaMag. The sediments were washed with lysis buffer, and then resuspended and heat-denatured in 2x Laemmli buffer (supplemented with 5% β-Mercaptoethanol). The final samples were run into SDS-PAGE gels before: 1) being transferred to membranes for Western Blotting; or 2) being sent to the proteomics core facility for mass spectrometry analysis. Anti-α-Adaptin 1/2 Antibody (C-8) (Santa Cruz Biotechnology,

Catalog #sc-17771) was used in WB to detect AP2 enrichments.

### Mass Spectrometry Analysis

Samples were digested overnight with trypsin (Pierce) following reduction and alkylation with DTT and iodoacetamide (Sigma–Aldrich). Following solid-phase extraction cleanup with an Oasis HLB µelution plate (Waters), the resulting peptides were reconstituted in 10 µL of 2% (v/v) acetonitrile (ACN) and 0.1% trifluoroacetic acid in water. 2 µL of each sample were injected onto an Orbitrap Fusion Lumos mass spectrometer (Thermo Electron) coupled to an Ultimate 3000 RSLC-Nano liquid chromatography systems (Dionex). Samples were injected onto a 75 μm i.d., 75-cm long EasySpray column (Thermo), and eluted with a gradient from 0-28% buffer B over 90 min. Buffer A contained 2% (v/v) ACN and 0.1% formic acid in water, and buffer B contained 80% (v/v) ACN, 10% (v/v) trifluoroethanol, and 0.1% formic acid in water. The mass spectrometer operated in positive ion mode with a source voltage of 2.4 kV and an ion transfer tube temperature of 275 °C. MS scans were acquired at 120,000 resolution in the Orbitrap and up to 10 MS/MS spectra were obtained in the Orbitrap for each full spectrum acquired using higher-energy collisional dissociation (HCD) for ions with charges 2-7. Dynamic exclusion was set for 25 s after an ion was selected for fragmentation.

Raw MS data files were analyzed using Proteome Discoverer v2.4 (Thermo), with peptide identification performed using Sequest HT searching against the human reviewed protein database from UniProt. Fragment and precursor tolerances of 10 ppm and 0.6 Da were specified, and three missed cleavages were allowed. Carbamidomethylation of Cys was set as a fixed modification and oxidation of Met was set as a variable modification. The false-discovery rate (FDR) cutoff was 1% for all peptides. To generate the volcano plot, Tubulin Beta 6 was used for Normalization.

The data was uploaded to the MassIVE data repository with accession number MSV000095338.

### Protein purification and GST pull-down assay

GST in a pGEX-6P-1 vector, GST-AP2-α-AD in a pGEX-2T-1 vector, and GST-AP2-β-AD in a pGEX-4T-1 vector were transfected and expressed in BL21(DE3) separately, and then affinity purified using GSTrap HP column (Cytiva). The affinity-purified GST fusion proteins were applied to a HiLoad 16/600 Superdex 200 pg column (Cytiva). Target proteins were collected and concentrated using Amicon Ultra-15 10K Centrifugal filters (Sigma-Aldrich), and stored in 20 mM HEPES, 150 mM NaCl, 1mM TCEP, pH 7.4.

In GST pull-down assays, purified GST, GST-AP2-α-AD, and GST-AP2-β-AD were pre-bound to anti-GST beads (Biolinkedin, #L-2004). Subsequently, these beads were separately added into 0.5 mg cell lysate of ARPE-HPV eGFP-CCDC32(FL). The reactions were allowed to proceed by rotation at 4°C for 2h, and then the beads were spun down at 1000 g, 4°C for 3 min. The sediments were washed with lysis buffer, and then heat-denatured in 2x Laemmli buffer (supplemented with 5% β-Mercaptoethanol) before running into SDS-PAGE gels, which followed by Western Blotting analysis.

### Transferrin receptor (TfnR) uptake assay

Internalization of TfnR was quantified by in cell ELISA following established protocols described (Chen et al., 2020; Conner and Schmid, 2003; Reis et al., 2015; Srinivasan et al., 2018). Briefly, ∼15,000 cells were seeded on gelatin-coated 96-well plate overnight. Next day, cells were starved in 37°C-warm PBS^4+^ (1× PBS buffer plus 0.2% BSA, 1mM CaCl_2_, 1mM MgCl_2_, and 5mM D-glucose) for 30 min. After starvation, the cell medium was replaced with ice-cold PBS^4+^ containing 5 µg/ml HTR-D65 (anti-TfnR mAb) (Schmid and Smythe, 1991). In parallel, some cells were kept at 4°C for the measurement cell surface TfnR (denoted as S) and blank controls (denoted as B), while some were incubated in a 37°C water bath for the indicated times for the measurement of internalized TfnR (denoted as I). For B and I, acid wash (0.2 M acetic acid and 0.2 M NaCl, pH 2.3) was applied to remove surface-bound HTR-D65. After washing with cold PBS. All cells were fixed with 4% PFA (Electron Microscopy Sciences) diluted in PBS for 30 min at 37°C. Subsequently, the cells were permeabilized with 0.1% Triton X-100 and blocked with Q-PBS (PBS, 2% BSA, 0.1% lysine, and 0.01% saponin, pH 7.4) for 2h. Surface-bound and internalized HTR-D65 were detected with HRP Goat anti-Mouse IgG (H+L) (BioRad) and o-phenylenediamine dihydrochloride (OPD, Sigma-Aldrich). Well-to-well variation of cell numbers was accounted for by BCA assays.

### TIRF and Epi-TIRF microscopy

Cells for imaging were seeded on gelatin-coated glass bottom dishes 35 mm (ibidi, #81218-800) for ∼12h before data acquisition. Live cell imaging was conducted with a Nikon Eclipse Ti2 inverted microscope that was equipped with: 1) an Apo TIRF/100x 1.49 Oil objective; 2) a Prime Back Illuminated sCMOS Camera (Prime BSI, 6.5 x 6.5µm pixel size and 95% peak quantum efficiency; 3) a M-TIRF module for epifluorescence (Epi) acquisition; 4) an H-TIRF module for TIRF acquisition, where penetration depth was fixed to 80nm; 5) an Okolab Cage Incubator for maintaining 37°C and 5% CO_2_.

For time-lapse TIRF imaging, 451 consecutive images were acquired at a frame rate of 1 frame/s for single channel or 0.5 frame/s for dual channels. For time-lapse Epi-TIRF imaging, 451 consecutive Epi and TIRF images were acquired nearly simultaneously at a frame rate of 0.66 frame/s. Perfect Focus System (PFS) was applied during time-lapse imaging.

The acquired data was analyzed using cmeAnalysis (Aguet et al., 2013; Jaqaman et al., 2008) and DASC (Wang et al., 2020), which are available at: https://github.com/DanuserLab/cmeAnalysis. Detailed protocols are available via this link. Briefly, cmeAnalysis was used to track the lifetime and fluorescence of clathrin-coated structures, and then DASC was applied to unbiasedly classify *bona fide* CCPs vs. abortive coats. Next, the classified CCP tracks were used to calculate CCP invagination (Δz), as previously described (Wang et al., 2020). Tracks that overlap with others or deviate from the properties of a diffraction-limited particle were excluded from the analysis.

### Platinum Replica Electron Microscopy (PREM)

The “unroofing” technique that mechanically removes the upper cell plasma membrane while preserving the ventral plasma membrane with CCSs and other structures for PREM analyses was performed essentially as described previously (Yang et al., 2022). Briefly, ARPE19 cells treated with siCCDC32 or siControl on coverslips were quickly transferred into ice-cold PEM buffer (100 mM PIPES−KOH, pH 6.9, 1LmM MgCl2 and 1 mM EGTA) containing 2 µM unlabeled phalloidin (Sigma , #P2141) and 10 µM taxol (Sigma-Aldrich, #T7402) and unroofed by a brief (1s) ultrasonic burst from a 1/8-inch microprobe positioned at ∼45° angle ∼3 mm above the coverslip and operated by Misonix XL2020 Ultrasonic Processor at 17-20% of output power. After sonication, the coverslips were immediately fixed with 2% glutaraldehyde in 0.1 M Na-cacodylate buffer, pH 7.3 for at least 20 min at room temperature.

Sample processing for PREM was performed as described previously (Svitkina, 2022). In brief, glutaraldehyde-fixed cells were post-fixed by sequential treatment with 0.1% tannic acid and 0.2% uranyl acetate in water, critical-point dried, coated with platinum and carbon, and transferred onto EM grids for observation.

PREM samples were examined using JEM 1011 transmission electron microscope (JEOL USA, Peabody, MA) operated at 100 kV. Images were acquired by an ORIUS 832.10W CCD camera (Gatan, Warrendale, PA) and presented in inverted contrast.

Based on the degree of invagination, the shapes of CCSs were classified into: flat CCSs with no obvious invagination; dome-shaped CCSs that had a hemispherical or less invaginated shape with visible edges of the clathrin lattice; and spherical CCSs that had a round shape with the invisible edges of clathrin lattice in 2D projection images. In most cases, the shapes were obvious in 2D PREM images. In uncertain cases, the degree of CCS invagination was determined using images tilted at ±10–20 degrees. The area of CCSs were measured using ImageJ and used for the calculation of the CCS occupancy on the plasma membrane.

### Online supplemental material

Fig. S1 shows the western blotting results that indicate the expression levels of exogenous CCDC32. Fig. S2 shows the representative TIRFM images of ARPE-HPV cells that stably express mRuby-CLCa and eGFP. It also shows the cohort-averaged fluorescence intensity traces of CCPs and CCP-enriched eGFP. Videos 1 and 2 shows addition time-lapse TIRFM imaging data for Fig. 2. Videos are accelerated 25-fold. Fig. S3 shows that siRNA-mediated knockdown of CCDC32 did not affect AP2 protein levels. Fig. S4 shows the fractions of CCPs in lifetime cohorts. Fig. S5 shows additional data for Fig. 4. Fig. S6 shows the AlphaFold 3.0 prediction of CCDC32 monomer structure and CCDC32-AP2-α complex structure. Fig. S7 shows a collection of representative TIRFM images from the main text showing cells that stably express the same amount of eGFP-CCDC32(FL), eGFP, eGFP-CCDC32(Δ78-98), and eGFP-CCDC32(1-54). Table S1 shows additional data for Fig. 5. Table S2 shows a list of primers used in this study.

## Data availability

Original data are available from the corresponding authors upon request.

## Acknowledgements

We thank the UTSW proteomics core facility for their help with sample processing and analysis. We thank Justin Bi for PREM analysis. We thank Zhenyang Chen for active participation in protein preparation. This work is supported by the National Natural Science Foundation of China (Grant No. 32200564 to Z.C.), the Natural Science Foundation of Hunan Province, China (Grant No. 2024JJ2045), the National Institutes of Health grants R35 GM 140832 to T.S., and the National Institutes of Health grant GM73165 to S.L.S.

## Author contributions

Z. Chen, Z. Yang, and C. Yang designed the experiments. Z. Chen, Z. Yang, C. Yang, T. Svitkina and S.L. Schmid interpreted the results and wrote the manuscript with input from all authors. Z. Yang, Y. Li and Z. Huang performed the TIRF and Epi-TIRF microscope imaging and data analysis. Z. Yang performed the co-IP, GST pull-down assays and TfnR uptake assays. C. Yang performed the PREM experiments. P. Xu performed FACS sorting. L. Han drew the cartoon illustrations. J. Pak did the AlphaFold3 modeling.

## Supplementary Information

**Video 1:** Time-lapse TIRFM imaging of ARPE-HPV eGFP-CLCa cells that were treated with control siRNA. Images were obtained at 1 frame/s and collected for 7.5 min. Video is accelerated 25-fold.

**Video 2:** Time-lapse TIRFM imaging of ARPE-HPV eGFP-CLCa cells that were treated with CCDC32 siRNA. Images were obtained at 1 frame/s and collected for 7.5 min. Video is accelerated 25-fold.

---

**Figure S1:**
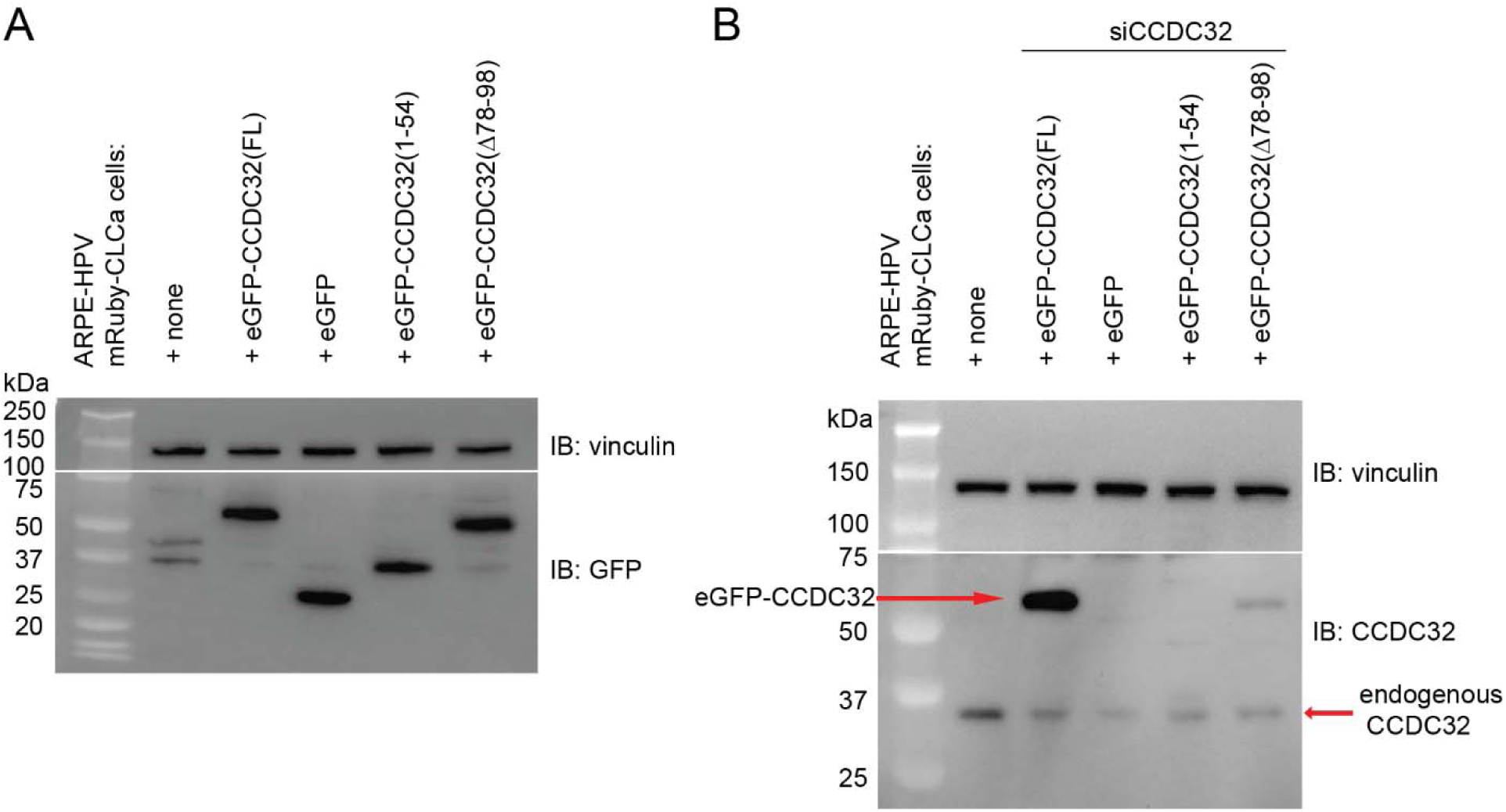
**(A)** Western blotting indicates similar expression levels of exogenous eGFP-CCDC32(FL), eGFP, eGFP-CCDC32(1-54) and eGFP-CCDC32(Δ78-98) in ARPE-HPV mRuby-CLCa cells. **(B)** siCCDC32 knocked down endogenous CCDC32 but not exogenously introduced eGFP-CCDC32(FL) from ARPE-HPV mRuby-CLCa cells. Note that the anti-CCDC32 antibody does not detect the eGFP-CCDC32(Δ78-98) as well as full-length and is unable to detect eGFP-CCDC32(1-54).

**Figure S2:**
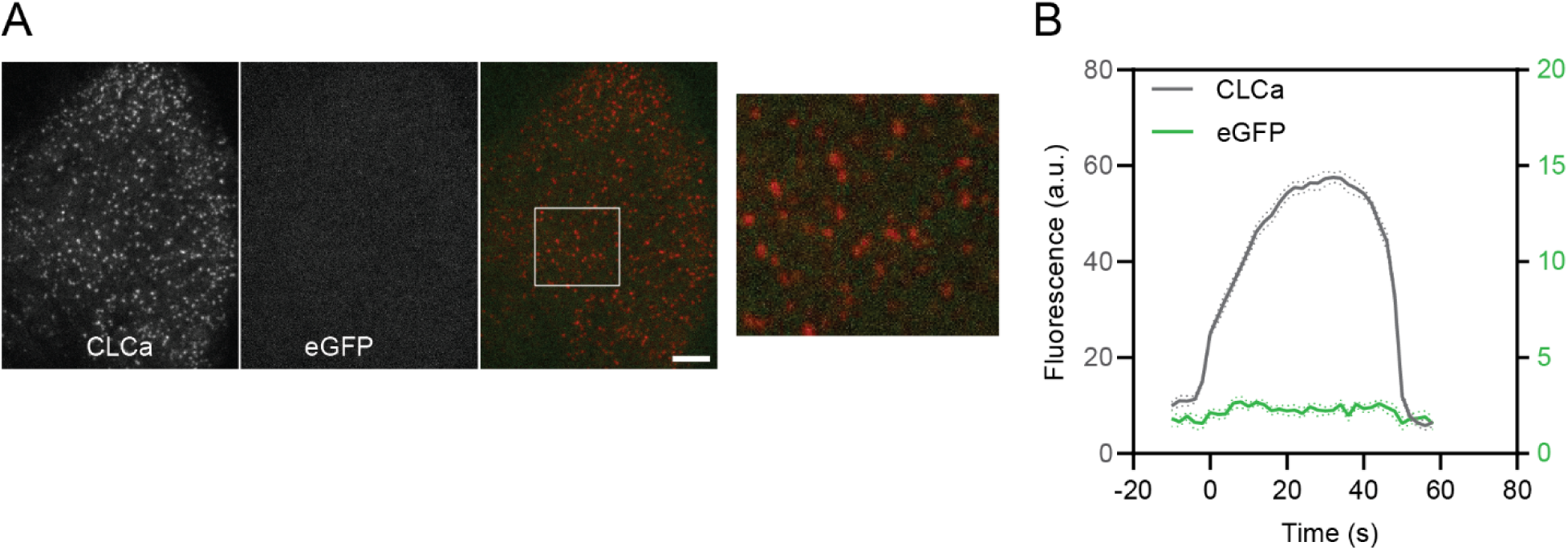
**(A)** Representative TIRFM images of ARPE-HPV cells that stably express mRuby-CLCa and EGFP. White ROI is magnified on the right. Scale bar = 5µm. **(B)** Cohort-averaged fluorescence intensity traces of CCPs and CCP-enriched EGFP. Number of tracks analyzed: 46833.

**Figure S3:**
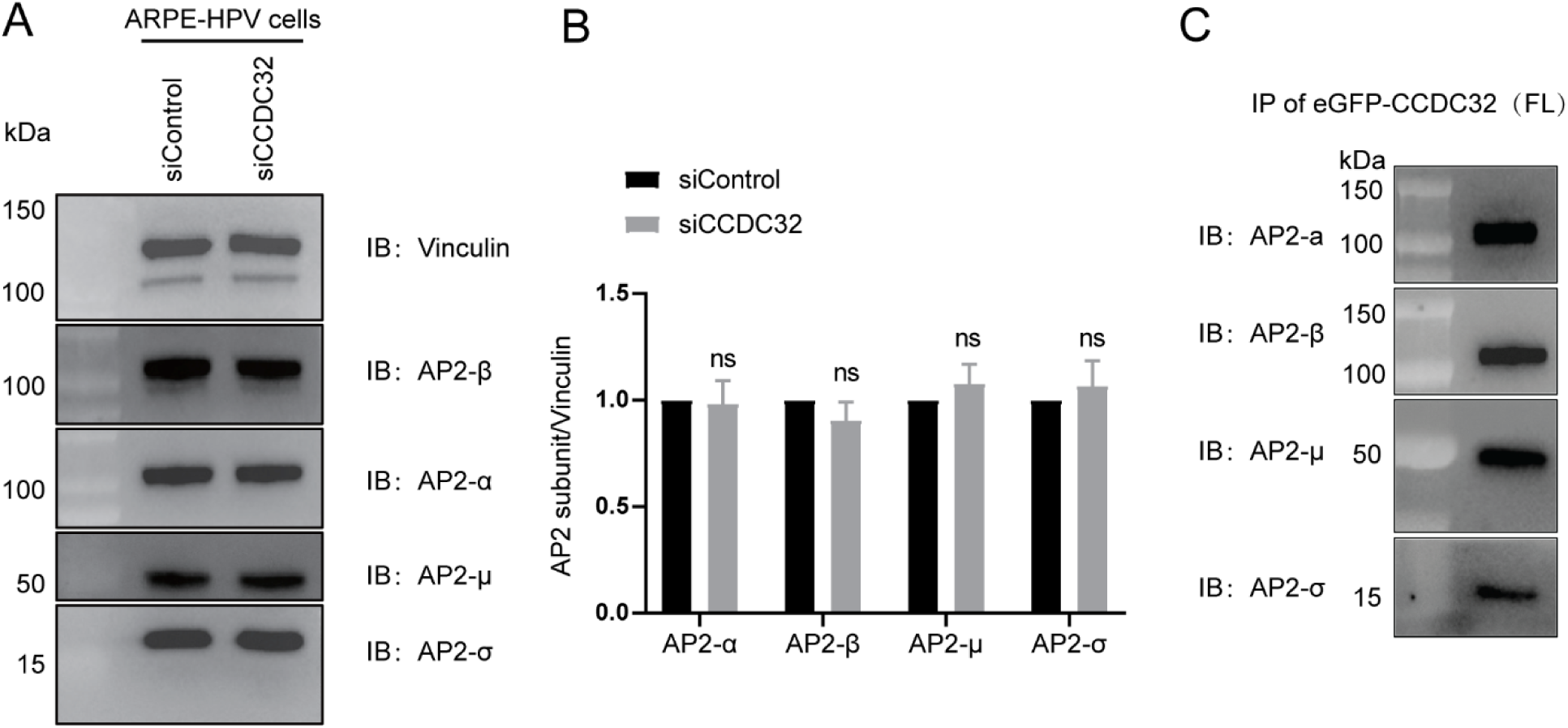
**(A-B)** siRNA-mediated knockdown of CCDC32 does not affect AP2 expression level. (A) Representative immunoblotting result of AP2 subunits from n=3 biological repeats. **(B)** Quantification of results from (A). Error bars indicate standard deviations. Statistical analysis is student’s t-test: ns, not significant. **(C)** All the four subunits of intact AP2 efficiently co-IP with eGFP-CCDC32(FL).

**Figure S4:**
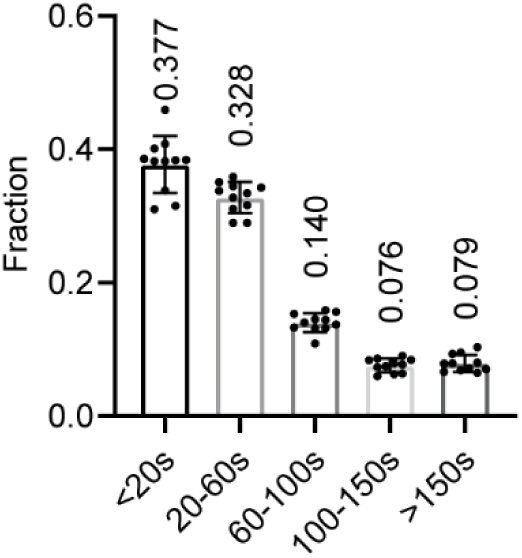
Fraction of CCPs in lifetime cohorts. Data was obtained from ARPE-HPV eGFP-CLCa cells that were treated with siCCDC32. The obtained data was processed with cmeAnalysis.

**Figure S5:**
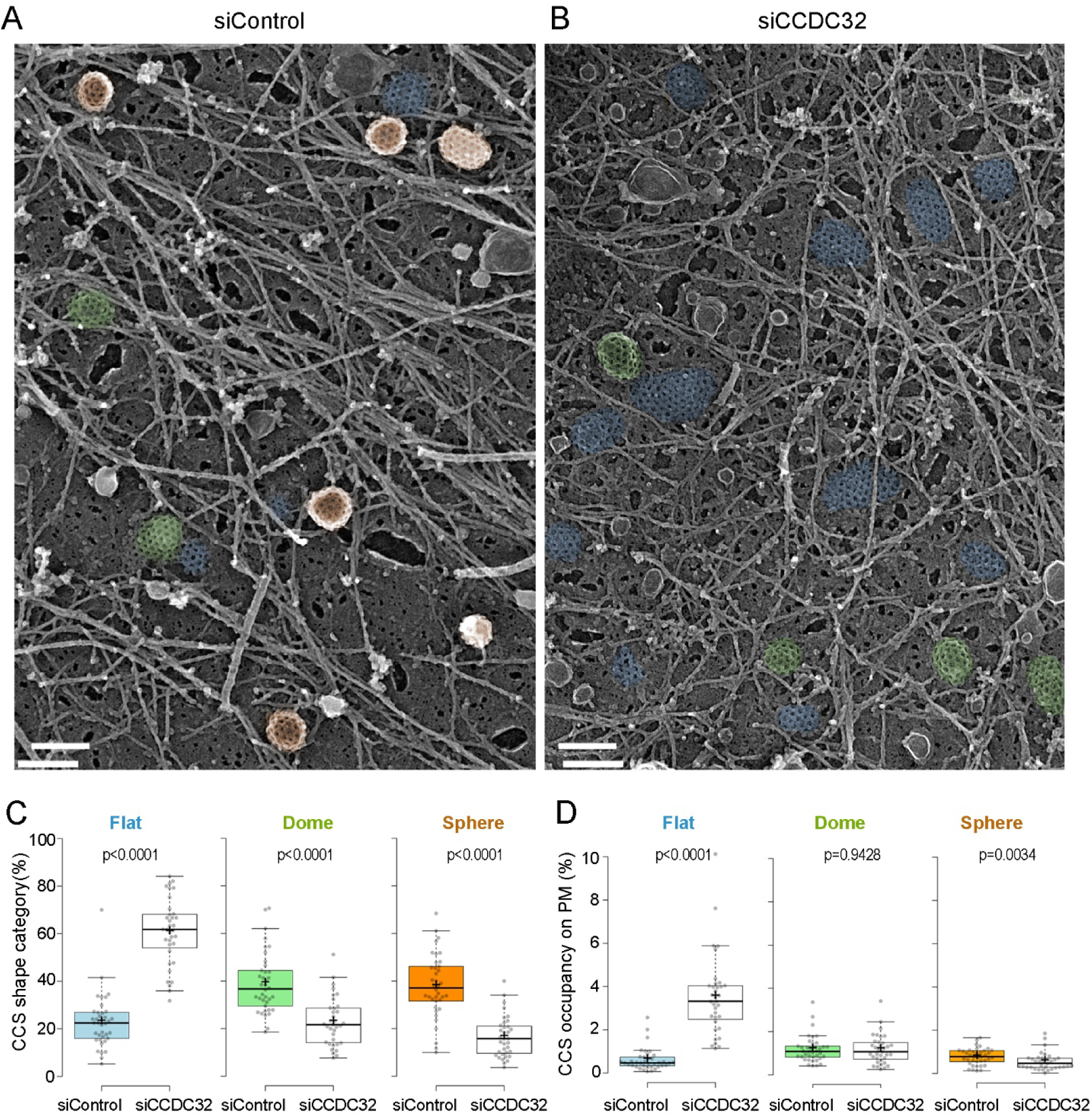
CCDC32 knockdown shifts the shapes of clathrin-coated structures. (A-B) Representative PREM images of ARPE-HPV cells treated with control siRNA (A) or CCDC32 siRNA (B). Scale bars = 200 nm. **(C-D)** Quantification of the CCS shape category (C) and CCS occupancy on PM (D). Each dot represents an individual fragment of the plasma membrane; the number of plasma membrane fragments analyzed is 38 for siControl and 35 for siCCDC32 from two independent experiments. Statistical tests were performed using either unpaired t-test or Mann-Whiney test depending on normality of distributed values.

**Figure S6:**
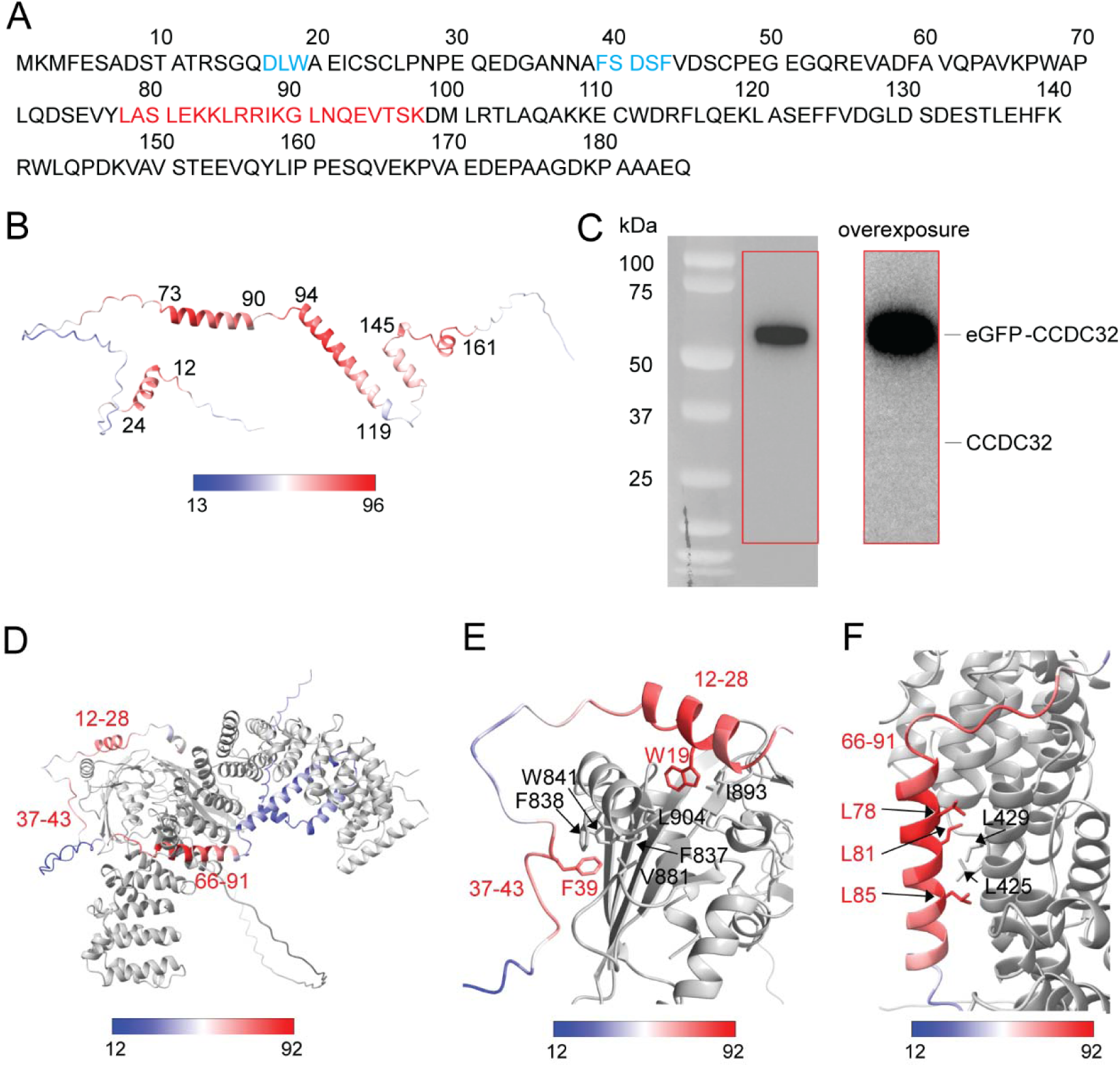
The sequence and structure of CCDC32. (**A**) Amino acid sequence of CCDC32. Blue font: canonical AP2 binding motifs; red font: predicted coiled-coil domain (aa78-98). (**B**) AlphaFold 3.0 modelling of full-length CCDC32 (Jumper et al., 2021; Varadi et al., 2021). Residues with pLDDT > 0.8 are labelled in red. **(C)** co-IP of eGFP-CCDC32 did not pulldown endogenous CCDC32. Red-cycled area was overexposed on the right. **(D-E)** AlphaFold 3.0 modelling of CCDC32 interaction with AP2-α (gene: AP2A2). For all panels, CCDC32 is colored by pLDDT and AP2-α is grey. (D) CCDC32-AP2A2 complex. CCDC32 residues with pLDDT > 0.7 are labelled in red. (E-F) Close up view of CCDC32-AP2A2 complex near CCDC32 residues (E) 12-28, 37-43 and (F) 66-91. Hydrophobic CCDC32 and AP2-α residues at the interface are labelled in red and black, respectively. Two canonical α-AD binding motifs in CCDC32 (^17^DLW^19^ and ^39^FSDSF^43^ can be docked on the AP2 appendage domain with high confidence; aa78-98 docks with high confidence to an alpha helix in the α-subunit encoded by aa418-438. Alphafold 3.0 modelling was performed using Alphafold Server (Abramson et al., 2024) and structure images were generated using ChimeraX (Meng et al., 2023).

**Figure S7:**
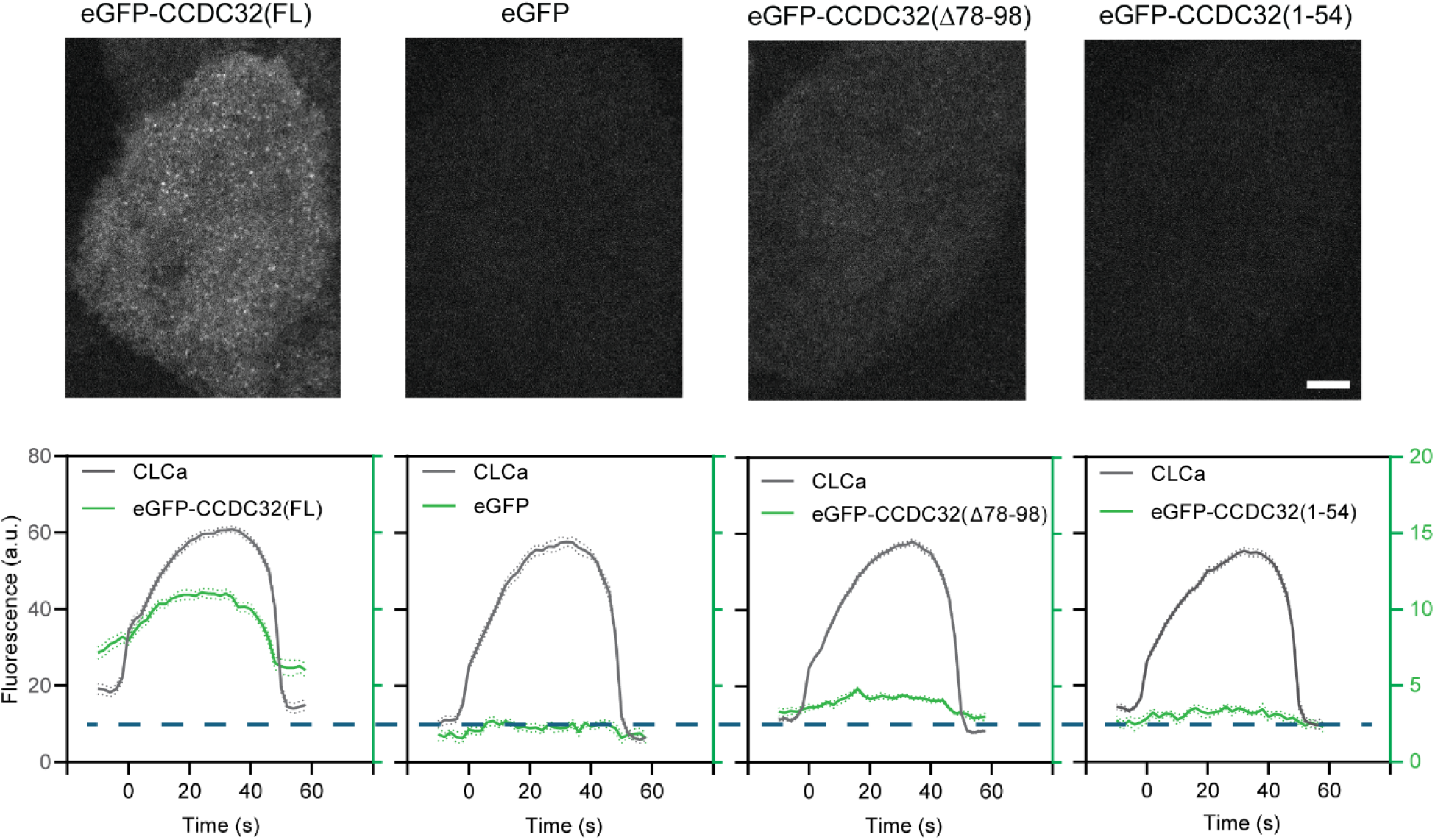
A collection of representative TIRFM images from the main text showing cells that stably express the same amount of eGFP-CCDC32(FL), eGFP, eGFP-CCDC32(Δ78-98), and eGFP-CCDC32(1-54). Diffuse PM fluorescence is greatest for eGFP-CCDC32(FL), diminished, but detectable for eGFP-CCDC32(Δ78-98), but absent for eGFP-CCDC32(1-54) and control eGFP. Scale bar = 5µm. Blue dashed line indicates the background PM fluorescence of eGFP.

**Table S1:** ID and abundances of key proteins detected in IP samples by Mass Spectroscopy analysis. #1-#3: 3 independent repeats of control IPs; #4-#6: 3 independent repeats of AP2-α-eGFP IPs.

| Gene | Abundance |  |  |  |  |  |
| --- | --- | --- | --- | --- | --- | --- |
| | Control IPs | | | AP2-α-eGFP IPs | | |
|  | #1 | #2 | #3 | #4 | #5 | #6 |
| AP2A1 | 2.E+06 | 1.E+06 | 6.E+05 | 1.E+08 | 1.E+08 | 2.E+08 |
| AP2A2 | 4.E+06 | 1.E+06 | 7.E+05 | 3.E+08 | 3.E+08 | 3.E+08 |
| AP2B1 | 7.E+06 | 6.E+06 | 3.E+06 | 7.E+08 | 6.E+08 | 6.E+08 |
| AP2M1 | 1.E+06 | 8.E+05 | 2.E+05 | 2.E+08 | 1.E+08 | 2.E+08 |
| AP2S1 | 1.E+05 | 5.E+04 | 9.E+03 | 2.E+07 | 2.E+07 | 5.E+07 |
| CCDC32 | 0.E+00 | 0.E+00 | 0.E+00 | 3.E+06 | 2.E+06 | 2.E+06 |
| AAK1 | 7.E+04 | 1.E+05 | 3.E+04 | 2.E+07 | 1.E+07 | 2.E+07 |
| EPS15 | 2.E+06 | 3.E+05 | 2.E+05 | 3.E+07 | 3.E+07 | 3.E+07 |
| EPS15L | 3.E+05 | 1.E+04 | 2.E+04 | 1.E+07 | 1.E+07 | 1.E+07 |
| FCHO2 | 4.E+04 | 3.E+04 | 2.E+04 | 1.E+06 | 1.E+06 | 1.E+06 |
| ITSN1 | 1.E+04 | 0.E+00 | 0.E+00 | 1.E+06 | 9.E+05 | 7.E+05 |
| NECAP1 | 9.E+04 | 8.E+04 | 3.E+04 | 9.E+06 | 7.E+06 | 1.E+07 |
| NECAP2 | 1.E+05 | 7.E+03 | 1.E+04 | 2.E+07 | 2.E+07 | 2.E+07 |
| AAGAB | 9.E+00 | 0.E+00 | 0.E+00 | 3.E+06 | 4.E+06 | 7.E+06 |

**Table S2:**
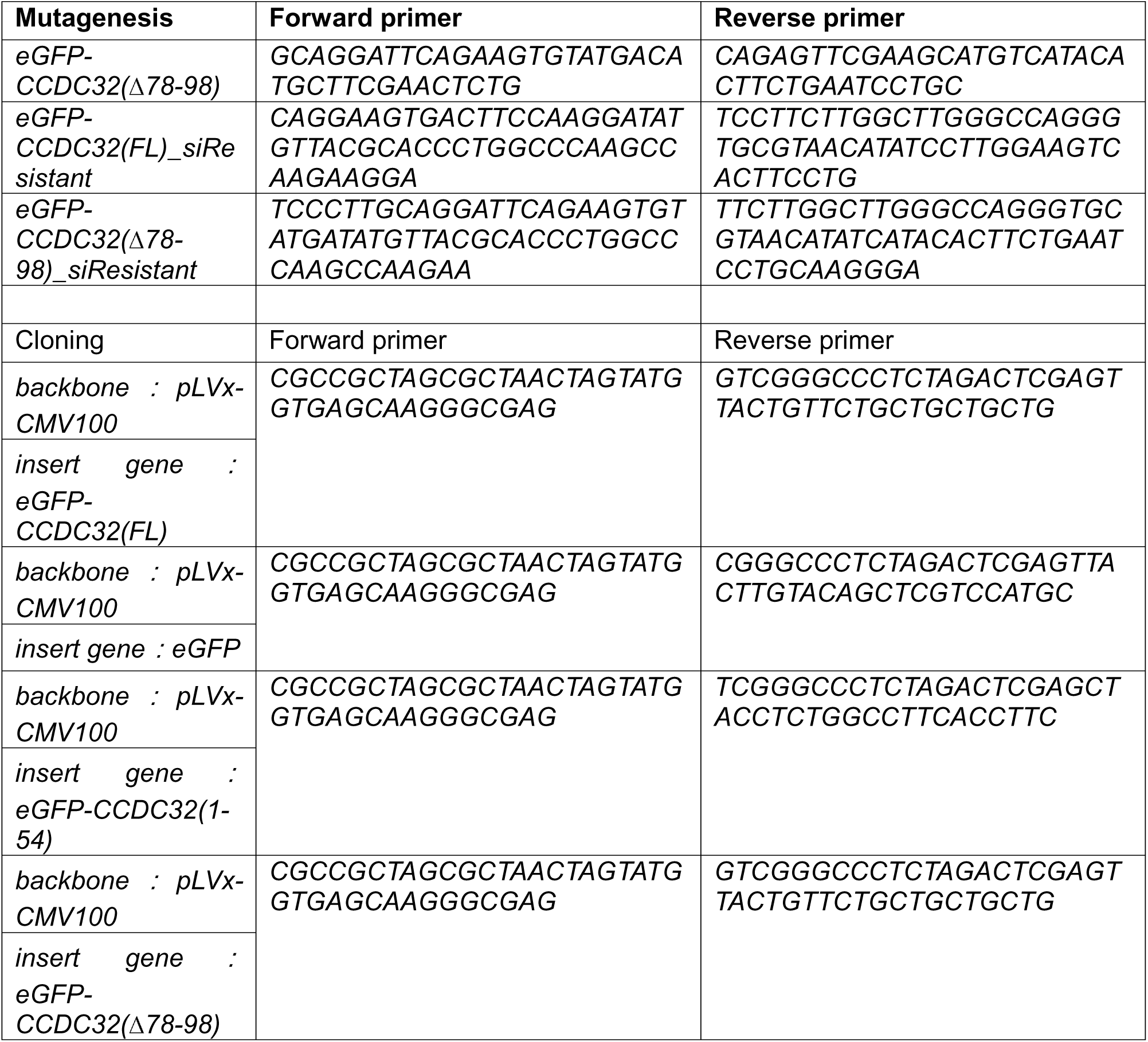
List of primers.

